# Multiple Holins Contribute to Extracellular DNA Release in *Pseudomonas aeruginosa* Biofilms

**DOI:** 10.1101/2020.10.28.358747

**Authors:** Amelia L. Hynen, James J. Lazenby, George M. Savva, Laura C. McCaughey, Lynne Turnbull, Laura M. Nolan, Cynthia B. Whitchurch

**Author notes:** These authors contributed equally (Laura M. Nolan, Cynthia B. Whitchurch).

## Abstract

Bacterial biofilms are comprised of aggregates of cells encased within a matrix of extracellular polymeric substances (EPS). One key EPS component is extracellular DNA (eDNA), which acts as a ‘glue’, facilitating cell-cell and cell-substratum interactions. We have previously demonstrated that eDNA is produced in *Pseudomonas aeruginosa* biofilms via explosive cell lysis. This phenomenon involves a subset of the bacterial population explosively lysing, due to peptidoglycan degradation by the endolysin Lys. Here we demonstrate that in *P. aeruginosa* three holins, AlpB, CidA and Hol, are involved in Lys-mediated eDNA release within both submerged (hydrated) and interstitial (actively expanding) biofilms, albeit to different extents, depending upon the type of biofilm and the stage of biofilm development. We also demonstrate that eDNA release events determine the sites at which cells begin to cluster to initiate microcolony formation during the early stages of submerged biofilm development. Furthermore, our results show that sustained release of eDNA is required for cell cluster consolidation and subsequent microcolony development in submerged biofilms. Overall, this study adds to our understanding of how eDNA release is controlled temporally and spatially within *P. aeruginosa* biofilms.

## Introduction

The biofilm mode of growth, in which bacterial aggregates are encased in a matrix of Extracellular Polymeric Substances (EPS), confers many advantages to bacteria, including increased resistance to antibiotics, predators, host cells and mechanical removal (1). The extreme difficulty in eradicating bacterial biofilms has important ramifications for industry and healthcare, where biofouling and medical-device associated biofilms cost billions of dollars each year and increase the morbidity and mortality of affected individuals (2–4). The complex EPS component of biofilms, which makes up to 90% of the biofilm biomass, consists of polysaccharides, extracellular DNA (eDNA), proteins, lipids and membrane vesicles (5). Together these components give the biofilm structure, stability and protection. Although individual biofilm components have been widely studied in many species (5–11), there is little understanding of the temporal and spatial production of each component and their roles in different stages of biofilm development. To develop means of preventing biofilm formation and disrupting mature biofilms, the complex interplay between cells, matrix components and the environment needs to be fully elucidated.

*Pseudomonas aeruginosa* is a WHO multi-drug resistant “priority pathogen” and is a model organism for studying biofilm structure and development (12). eDNA is most notably described as having ‘glue’ like properties, facilitating cell-cell and cell-substratum interactions in both submerged (hydrated) and interstitial (actively expanding) biofilms (8,13,14). Other roles of eDNA in the biofilm matrix include coordinating migration, facilitating horizontal gene transfer and altering antibiotic tolerance (14–18). We previously demonstrated the requirement of eDNA for biofilm formation by *P. aeruginosa* in submerged (13) and interstitial biofilms (14) using the DNA degrading enzyme Deoxyribonuclease I (DNase I) and that eDNA in both interstitial and the early developmental stages of submerged *P. aeruginosa* biofilms is produced by explosive cell lysis (19). Explosive cell lysis involves a subset of the bacterial population rapidly transitioning from a rod-shaped cell to a round morphotype, due to peptidoglycan degradation by the endolysin Lys. This dramatic and rapid event results in expulsion of the cellular contents and membrane vesicles into the biofilm milieu.

Lys (PA0629) is a bacteriophage related endolysin encoded in the R- and F-pyocin gene cluster, along with the holin Hol, which are both required for pyocin release from producing cells (20). In *P. aeruginosa,* overexpression of Hol has been previously shown to increase lysis (21). Holin proteins form pores in the inner membrane to transport endolysins to the periplasm and are usually associated with an antiholin that protects cells from their activity (22). Together the holin/antiholin system acts as a gatekeeper to the activity of Lys. Disruption of the outer membrane is also required for cell lysis and this is usually achieved by proteins called spannins (23). Lys is the only putative endolysin encoded on the *P. aeruginosa* PAO1 genome, however, Hol is not the only holin. Two other putative holins have been previously reported, namely, AlpB and CidAB. AlpB is part of a DNA-damage dependent self-lysis pathway in *P. aeruginosa,* which has been linked to eDNA release in broth cultures and increased colonization of the murine lung (24). The Cid (and associated Lrg) system shares similarities with the Bax/Bcl-2 family of proteins that are involved in the regulation of apoptosis in eukaryotes (25). In *Staphyloccus aureus* CidA is involved in eDNA release (26,27). In *P. aeruginosa* CidA is the putative holin that is homologous to CidA holins in other bacteria. In *P. aeruginosa* a double mutant of *cidA* as well as the uncharacterized downstream gene *cidB* (PAO1Δ*cidAB*) have been linked to cell death and the dispersal stage of submerged biofilm development (11). Although both of the holins AlpB and CidA have been associated with programmed cell death (PCD) pathways and eDNA release, no associated endolysins have been identified for these systems.

In our previous study we hypothesised that Hol is responsible for transportation of Lys across the inner membrane, and subsequent explosive cell lysis, as it is found alongside Lys in the pyocin gene cluster (19). In the current study we show that the three holins, Hol, AlpB and CidA, are involved in Lys-mediated explosive cell lysis. We demonstrate that each holin plays a different role in this process depending upon the type of biofilm (actively expanding or submerged) as well as the stage of submerged biofilm development. Furthermore, within submerged biofilms we show that the sites of explosive cell lysis-mediated eDNA release events correspond to the sites of subsequent cell cluster formation in the early stages of submerged biofilm formation, prior to microcolony formation. Additionally, we show that continual release of eDNA within these cell clusters is required for cell cluster and microcolony consolidation. To our knowledge this is the first report demonstrating that eDNA is required for initiating cell clustering and microcolony formation. Overall, this work establishes a link between the activity of *P. aeruginosa* holin-mediated explosive cell lysis resulting in eDNA release and the spatial regulation of biofilm formation and consolidation.

## Methods

### Bacterial strains and plasmids

*P. aeruginosa* and *E. coli* strains used in this study, as well as plasmids, are listed in Table S1. The primers used in this study are listed in Table S2. In-frame deletions of *hol* and *alpB* were constructed in *P. aeruginosa* strain PAO1 by allelic exchange using the Flp-FRT recombination system for site specific excision of chromosomal sequences as previously described (28). Briefly, 1 kb sections that flanked the *hol* and *alpB* regions were synthesized by GeneArt Gene Synthesis (Thermo Scientific), creating pALH3 and pALH9, respectively. This also included 100 bp of the 5’ and 3’ ends of the target gene with a *Hind*III site introduced in the middle. The *FRT-GmR* cassette from pPS856 (29) was subcloned into the internal *Hind*III site of pALH3 and pALH9 resulting in pALH5 and pALH10 containing the flanking regions of *hol* and *alpB* separated by the *FRT-GmR* cassette. The resultant clones were then digested with *Spe*I and cloned into the suicide vector pRIC380, resulting in pALH7 and pALH11. The pRIC380 vector contains the genes *sacB*/*sacR*, which results in sensitivity to sucrose and *oriT* which enables conjugal transfer. The resultant clones were transformed into the *E. coli* donor strain S17-1 in preparation for conjugal transfer into *P. aeruginosa* PAO1 strains. The *GmR* gene was then excised using the pFLP2 plasmid that expresses the Flp recombinase as described previously (28) creating *P. aeruginosa* strains with the *hol* or *alpB* regions deleted and replaced with an FRT sequence. Allelic exchange mutants were confirmed by PCR of isolated chromosomal DNA. Complementing plasmids were constructed by cloning the wild type gene synthesized by GeneArt Gene Synthesis (Thermo Scientific) into the multiple cloning site of pJN105 using *Spe*I and *Sac*I restriction enzymes from New England Biolabs (Australia). To create the double and triple mutants, a step-wise deletion of each holin was accomplished, by following the same procedure as described above, starting with the PAO1Δ*cidAB* strain (11) (double and triple mutants) or the PAO1Δ*hol* strain (double mutant).

### Growth conditions

All *P. aeruginosa* strains were cultured in lysogeny broth (LB) or cation adjusted Mueller Hinton broth (CAMHB) and cultured at 37°C overnight at 250 rpm, or on LB agar (1.5% (w/v)) and incubated at 37°C overnight. *E. coli* strains were cultured in LB overnight at 37°C and 250 rpm, or on LB agar (1.5% (w/v)) and incubated at 37°C overnight. Media was supplemented where appropriate with antibiotics at the following concentrations: For *E. coli* 100 μg mL^−1^ ampicillin (Astral Scientific), 50 μg mL^−1^ kanamycin sulphate (Astral Scientific) and 10 μg mL^−1^ gentamicin sulphate (Sigma-Aldrich) and for *P. aeruginosa* 50 μg mL^−1^ gentamicin sulphate . L-Arabinose (Sigma-Aldrich) was added at 0.02% (w/v) to induce gene expression under the control of *araBAD* promoter on pJN105 in *P. aeruginosa*.

### Interstitial biofilm assays and microscopic analysis

Interstitial biofilm assays were performed as described previously (14). Briefly, gellan gum-solidified nutrient media (TMGG; 0.4 × LB, 0.1% MgSO_4_7H_2_O (w/v), 0.8% (w/v) GelGro gellan gum (MP Biomedicals)), was evenly spread across sterilized microscope slides. Fluorescent stains (TOTO-1 iodide; 1 μM, Life Technologies, and ethidium homodimer-2 (EthHD-2); 1 μM, Biotium) were added to the molten media immediately prior to pouring where indicated. Once set, a small inoculum of the strain of interest was applied to the TMGG-coated slide, a coverslip (0.13-0.16 mm thick; Menzel Glaser) placed on top and the slide incubated at 37°C in a humidified chamber for 4 h. The slides were imaged on a Nikon Ti inverted research microscope with a × 100 1.45 numerical aperture (NA) PlanApo objective, using NIS Elements acquisition software (Nikon Instruments, Tokyo, Japan), solid state illumination (Lumencor, Beaverton, OR, USA), Cascade 1Kx1K EMCCD camera (Photometrics) and fitted with an environmental chamber (ClearState Solutions, Mt Waverley, VIC, Australia). For quantitative analysis of eDNA release sites in interstitial biofilms, series of overlapping images spanning the outermost leading edge through to the inoculation site were obtained and stitched using the NIS Elements acquisition software (Nikon Instruments, Tokyo, Japan). The area analysed was 148.75 μm × 400.80 μm (59.6 mm^2^) back from the leading edge of the interstitial biofilm, which contains a monolayer of actively migrating cells and excludes older cells with compromised membranes at the inoculation site (Figure S1, S2). Within this area the number of eDNA release sites and the area covered by cells within the biofilm was quantified using FIJI (30). Briefly, the ‘Subtract Background’ process (rolling ball radius = 50 pixels) was applied to both the phase contrast and fluorescent images followed by ‘Auto-Thresholding’ using a ‘Triangle’ method with a dark background. The ‘Analyze Particles’ process was used to calculate the area of each eDNA release events (5-Infinity μm^2^ to only include explosive eDNA release events; Circularity 0.00-1.00) and the ‘Measure’ process was used to calculate the area covered by cells within the biofilm.

### Submerged biofilm assays and microscopic analysis

Submerged biofilm assays were performed as described previously (19). To avoid carryover of eDNA from the overnight cultures, cells were washed three times with fresh CAMHB, diluted 1/100 in CAMHB, cultured at 37°C for 2 h (250 rpm), diluted again 1/100 in CAMHB, 300 μL transferred to an μ-Slide 8 well ibiTreat microscopy chamber (ibidi, GmbH, Germany) and then incubated statically at 37°C for the indicated time. For submerged biofilm assays with complementation plasmid pJN105 L-arabinose (0.02 % (w/v)) was also added to the inoculum.

To visualise biofilm formation after 8 h of static culture, wells were washed twice with fresh media before CAMHB containing the eDNA stain (EthHD-2 (1 μM; Biotium)) was added to the wells. Biofilms and cells at the substratum were then imaged with phase contrast and wide-field fluorescence microscopy (Olympus IX71, × 40 objective). The size of microcolonies formed and raw integrated density after 8 h was determined using FIJI (30). Briefly, for phase contrast images the ‘Find Edges’ and ‘Sharpen’ processes were performed, followed by a ‘Gaussian Blur’ filter (Sigma (Radius) = 2.00) and ‘Auto-Thresholding’ using a ‘Triangle’ method with a dark background. The ‘Analyze Particles’ process was used to calculate the size of each microcolony (90-Infinity μm^2^; Circularity 0.00-1.00). The ROI for each microcolony in a field of view was overlaid onto the corresponding fluorescent image which had been processed using ‘Auto-Thresholding’ with a ‘Triangle’ method and dark background. The ‘Analyze Particles’ process was used to calculate the Raw Integrated Density of the fluorescent signal within the bounds of overlaid ROI (30-Infinity μm^2^; Circularity 0.00-1.00).

To observe eDNA release and cell cluster formation during the development of the submerged biofilms cells were incubated as described for the 8 h static culture setup, except in this case TOTO-1 iodide (1 μM; Life Technologies) was added with the inoculum to each well of the microscopy chamber. Time-lapse imaging (phase contrast and wide-field fluorescence microscopy) commenced at 1 hr post inoculation and continued at 30 min intervals for 5.5 h total (Nikon Ti inverted research microscope, × 100 objective). eDNA release events and cell cluster formation during the development of submerged biofilms were analyzed using FIJI (30). The same processes as described above for the 8 h microcolonies were used to generate a binary fluorescent image. The ‘Analyze Particle’s process was used to determine the Raw Integrated Density of each eDNA release event over time (0-Infinity μm^2^; Circularity 0.00-1.00). Raw Integrated Density values greater than 100,000 were included in further analysis. The time of cell cluster formation and the X, Y coordinate were determined manually using the ‘Multi-Point’ tool in FIJI.

### Statistical analysis

Effects of gene deletions were typically estimated using linear mixed models, with deletions included as fixed effects and field of view nested within replicates as random effects. Models were estimated using the lme4 (version 1.1-23) (31) and lmerTest (version 3.1-2) (32) packages for R (version 4.0.1) (33). Effects on counts were estimated using negative binomial mixed models or Poisson regression where there was no evidence of over-dispersion; effects on continuous outcomes using linear models following log-transformation where necessary. See Supplementary Tables S3-S10 for effect sizes, 95% CIs and p-values associated with relevant Figures, and Figure captions for details of specific analyses.

## Results

### Hol and AlpB contribute to explosive cell lysis in interstitial biofilms

To identify the holin(s) involved in Lys-mediated explosive cell lysis during twitching motility-mediated biofilm expansion, we cultured interstitial biofilms of PAO1, PAO1Δ*alpB*, PAO1Δ*cidAB,* PAO1Δ*hol* and PAO1Δ*lys* in the presence of the membrane impermeant nucleic acid stain EthHD-2 and quantified the number of explosive cell lysis-mediated eDNA release sites across 30 random fields of view. We observed a large number of cells taking up EthHD-2 stain in the inoculum (Figure S1), likely because these older cells have compromised membranes. To avoid the contribution of dead cells in our analyses we quantified eDNA events only within the area at the leading edge of the interstitial biofilm that was comprised of a monolayer of actively migrating cells (Figure S1, S2). In this area we observed numerous sites of eDNA release events for PAO1 (Figure 1A, C, Figure S1). As described previously (19), we observed very few eDNA release events/area for PAO1Δ*lys* (rate ratio=0.04, 95% CI=0.02-0.08, p<0.001; Table S3) (Figure 1B, C, Figure S2). We did not see a significant difference in the rate of eDNA events/area for PAO1Δ*alpB*, PAO1Δ*cidAB* or PAO1Δ*hol* compared to PAO1 (p=0.702, 0.705 or 0.768, respectively; Table S3) (Figure 1A, C, Figure S1-2).

**Figure 1.**
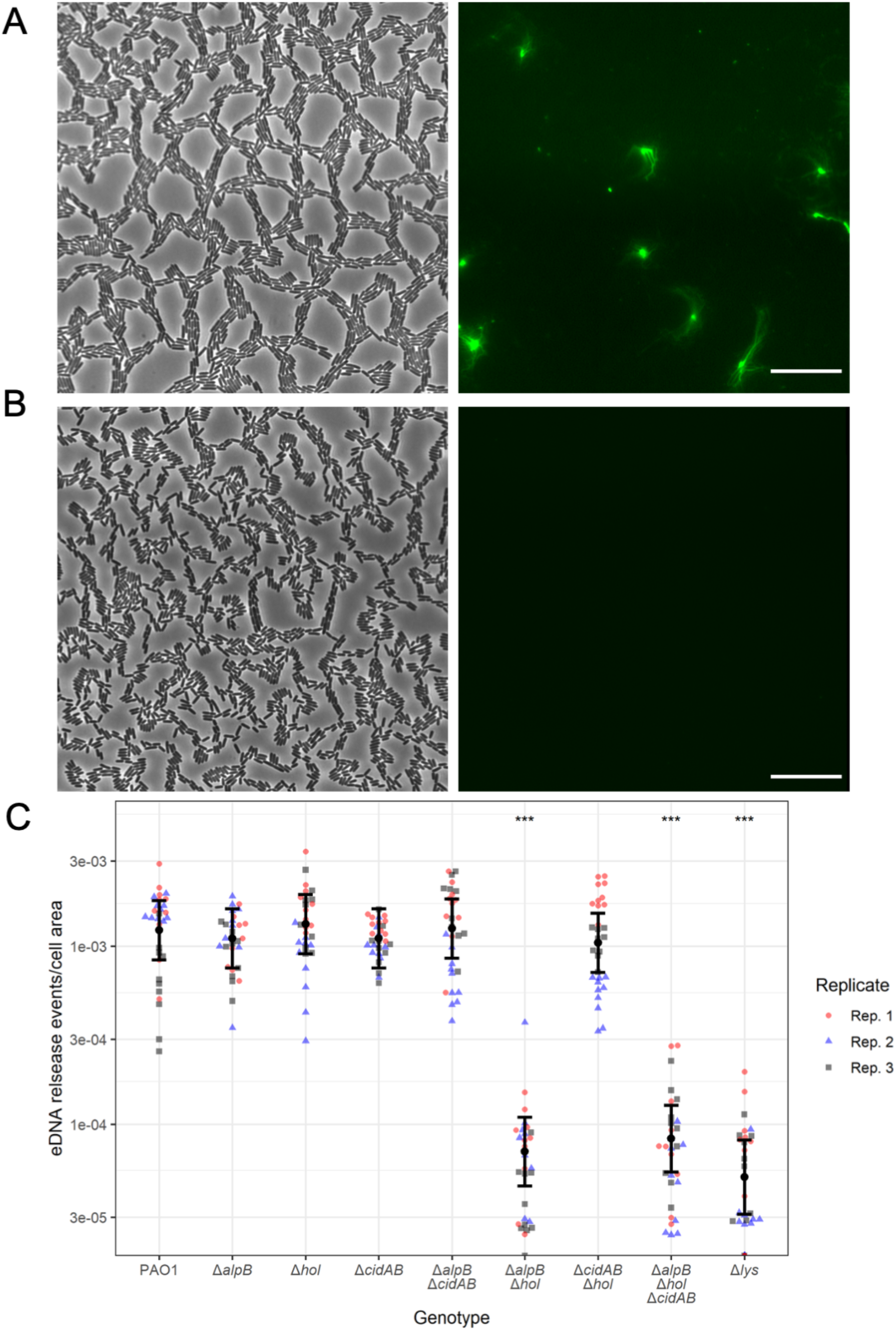
Hol and AlpB are both required for Lys-mediated eDNA release in interstitial biofilms. Interstitial biofilms of PAO1, PAO1Δ*alpB*, PAO1Δ*cidAB*, PAO1Δ*hol*, PAO1Δ*alpB*Δ*cidAB*, PAO1Δ*alpB*Δ*hol*, PAO1Δ*cidAB*Δ*hol*, PAO1Δ*alpB*Δ*cidAB*Δ*hol* and PAO1Δ*lys* were allowed to expand for 4 h at 37°C prior to imaging. (A-B) Representative phase-contrast (left) or EtHD-2 stained eDNA (green, right) images for (A) PAO1 or (B) PAO1Δ*alpB*Δ*cidAB*Δ*hol.* Images of PAO1Δ*alpB*, PAO1Δ*cidAB*, PAO1Δ*hol*, PAO1Δ*alpB*Δ*cidAB* and PAO1Δ*cidAB*Δ*hol* were indistinguishable from (A) and PAO1Δ*lys* and PAO1Δ*alpB*Δ*hol* were indistinguishable from (B). Scale=20 μm. (C) Numbers of eDNA release sites per area of cells within each field of view in the expanded biofilm area. The estimated means with 95% CIs are presented, which are from analysis of 10 individual fields of view from 3 biological replicates (n=30). Means and CIs were estimated using a negative binomial mixed effects regression model, with genotype nested within biological replicate as random effect, and genotype as a fixed effect. *** p<0.001 compared to PAO1. All other comparisons to PAO1 are ns i.e. p>0.05. See Table S3 for estimates, 95% CIs and p-values.

To determine whether there was redundancy in the contribution of these holins or if there were additional holins that might be involved in Lys translocation to the periplasm, we created a triple holin unmarked deletion mutant (PAO1Δ*alpB*Δ*cidAB*Δ*hol*) and deletions of combinations of two holins (PAO1Δ*alpB*Δ*cidAB*, PAO1Δ*alpB*Δ*hol*, PAO1Δ*cidAB*Δ*hol*). The triple mutant had a similar rate of explosive cell lysis events in interstitial biofilms compared to PAO1Δ*lys* demonstrating that there are likely to be no additional holins in PAO1 (rate ratio=0.07, 95% CI=0.04-0.12, p<0.001; Table S3) (Figure 1B, C, Figure S2). Neither PAO1Δ*alpB*Δ*cidAB* nor PAO1Δ*cidAB*Δ*hol* had a significant difference in the rate of eDNA release events compared to PAO1 (p=0.929 or p=0.556, respectively; Table S3) (Figure 1A, C, Figure S2). However, PAO1Δ*alpB*Δ*hol* was just as defective in explosive cell lysis events as the triple holin mutant and PAO1Δ*lys* (rate ratio=0.06, 95% CI=0.03-0.10, p<0.001; Table S3) (Figure 1B, C, Figure S2), suggesting that either AlpB or Hol are sufficient to facilitate Lys-mediated eDNA release in actively migrating interstitial biofilms. To our knowledge, this is the first description of multiple holins facilitating translocation of a single endolysin.

### Hol, AlpB and CidA mediate microcolony formation in submerged biofilms

We have previously demonstrated that eDNA is required for submerged biofilm development in *P. aeruginosa* (13) and that Lys-mediated explosive cell lysis is required for microcolony formation in submerged *P. aeruginosa* biofilms (19). Based on our findings that Hol and AlpB are required for explosive cell lysis in interstitial biofilms, we hypothesised that at least one of these holins and/or CidA would also be involved in microcolony formation in submerged biofilms. To investigate this, submerged biofilms of PAO1, PAO1Δ*alpB*, PAO1Δ*cidAB,* PAO1Δ*hol*, PAO1Δ*alpB*Δ*cidAB*Δ*hol* and PAO1Δ*lys* were cultured for 8 h, washed to remove unattached cells, then stained with the eDNA stain EthHD-2. The number of microcolonies, their size and the amount of eDNA present within microcolonies was measured in over 30 random fields of view for each strain. The amount of eDNA was determined by the integrated density value of the corresponding fluorescence signal within the microcolony.

In accordance with our previous observations (19), PAO1Δ*lys* was completely defective in producing eDNA through explosive cell lysis and did not form any microcolonies (Figure 2A). The triple holin deletion strain (PAO1Δ*alpB*Δ*cidAB*Δ*hol*) was just as defective as PAO1Δ*lys* in this assay (Figure 2A) again suggesting that there are no additional holins contributing to Lys translocation in PAO1. For PAO1, each field of view contained between 4-19 (mean of 8) microcolonies (Figure 2B). All PAO1 microcolonies were distinct and well-structured with eDNA present throughout the whole structure (Figure 2A). Their sizes ranged in area from 30-16,953 μm^2^, with a median area of 127 μm^2^ (Figure 2C). The size of the microcolony correlated to the amount of eDNA present (Figure S3).

**Figure 2.**
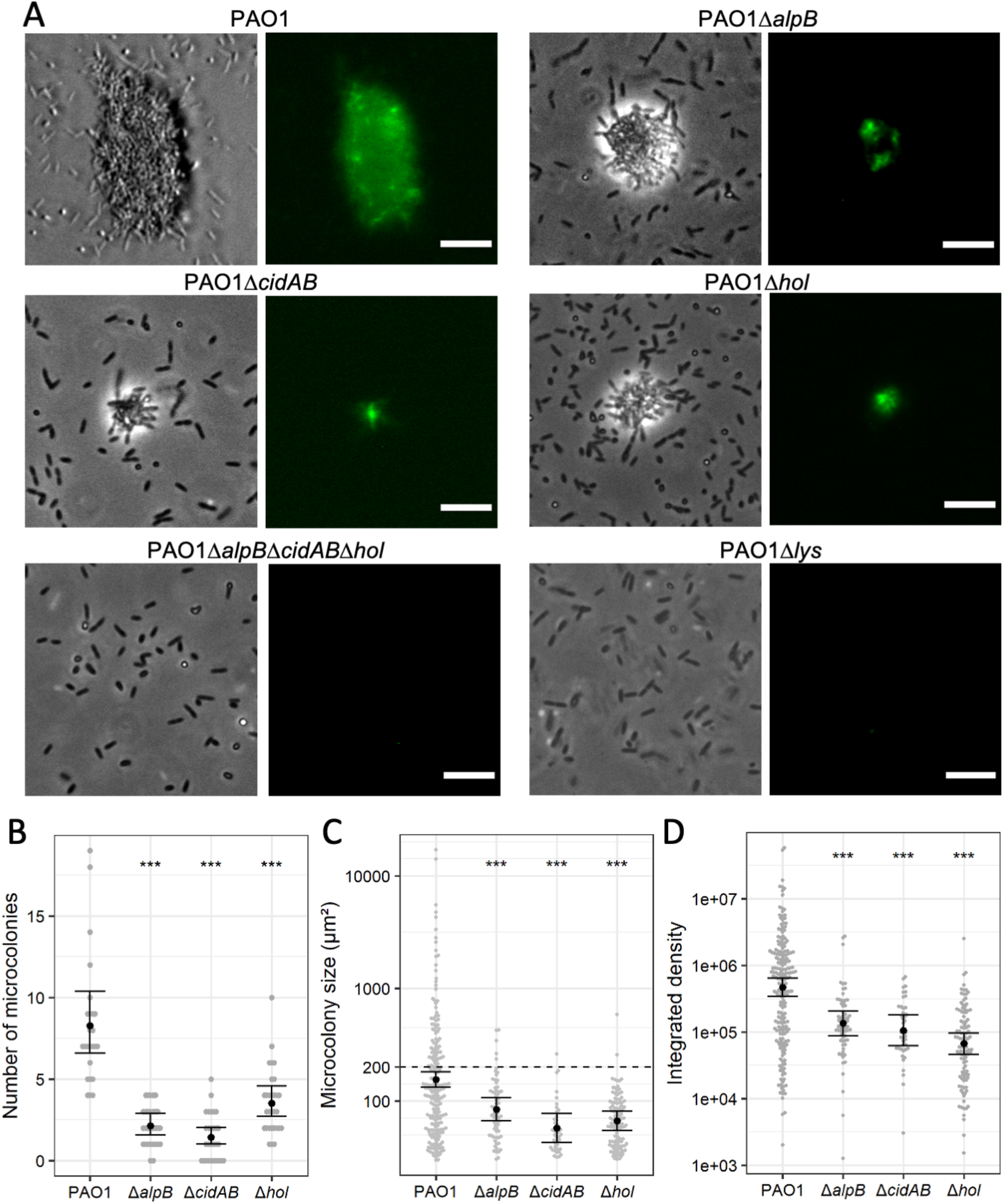
Mutations in *alpB, cidAB* and *hol* result in fewer microcolonies that are smaller and contain less eDNA than PAO1 under submerged biofilm conditions. Microcolonies formed by PAO1, PAO1Δ*alpB,* PAO1Δ*cidAB*, PAO1Δ*hol*, PAO1Δ*alpB*Δ*cidAB*Δ*hol* and PAO1Δ*lys* after 8 h incubation at 37°C. (A) Representative images of each strain with phase contrast (left) and eDNA (EthHD-2, right) are shown. Random fields of view were imaged for each strain across 3 biological replicates (n=25 (PAO1), n=30 (PAO1Δ*alpB,* PAO1Δ*hol*, PAO1Δ*alpB*Δ*cidAB*Δ*hol* and PAO1Δ*lys)* and n=29 (PAO1Δ*cidAB*)). Scale=10 μm. (B) Number of microcolonies, (C) microcolony size (μm^2^) or (D) integrated density of fluorescent signal (eDNA) in microcolonies. No microcolonies were observed in PAO1Δ*lys* and PAO1Δ*cidAB*Δ*hol*Δ*alpB.* For B-D data are presented as the mean ± SE; *** p<0.01, comparison of each deletion to PAO1. For (B) p-values and 95% CIs were calculated by Poisson regression comparing counts of microcolonies in each field of view over genotype. For (C) and (D) p-values were calculated by linear mixed models of log(outcome) on genotype, including a random effect of field of view. See Supplementary Table S4-6 for estimates, 95% CIs and p-values.

For each of the single deletion holin mutants we visualised significantly fewer microcolonies compared to PAO1, with between 0-4 (mean of 2.2) microcolonies per field of view for PAO1Δ*alpB,* between 0-5 (mean of 1.4) microcolonies per field of view for PAO1Δ*cidAB* and between 1-10 (mean of 3.6) microcolonies per field of view for PAO1Δ*hol* (Figure 2B; Table S4). Overall the microcolonies for the single holin mutants were significantly smaller than PAO1 ranging from 30-429 μm^2^ (mean of 77 μm^2^) for PAO1Δ*alpB,* 31-262 μm^2^ (mean of 51 μm^2^) for PAO1Δ*cidAB* and 30-586 μm^2^ (mean of 63 μm^2^) for PAO1Δ*hol* (Figure 2C; Tables S5). There was no evidence that the relationship between amount of eDNA (integrated density) and microcolony size varied between PAO1 and the single holin mutants (Figure S3). However, the microcolonies formed by the holin mutants contained significantly less eDNA and were not as tightly formed as the microcolony structures observed for PAO1 (Figure 2D; Table S6).

Wildtype microcolony formation was restored to each of the individual holin mutants by exogenous expression of the cognate holin gene, indicating that these defects in microcolony formation were due to loss of the holin and not due to secondary mutation (Figure S4; Table S7). Our data thus far demonstrates that Lys-mediated eDNA release facilitated by the holins AlpB, CidA and Hol is required for the formation of densely packed microcolonies in submerged biofilms imaged at 8 h and that these microcolonies always contain eDNA.

### eDNA release initiates the formation of cell clusters in submerged biofilms

Given the requirement for AlpB, CidA and Hol in the formation of microcolonies in submerged biofilms imaged at 8 h we next investigated the contribution of these holins to microcolony formation. We used phase contrast and fluorescence time lapse microscopy of PAO1, PAO1Δ*alpB,* PAO1Δ*cidAB* and PAO1Δ*hol* cultured in the presence of the membrane impermeant nucleic acid stain TOTO-1 from 1 h post inoculation and recorded the coordinates of eDNA release events and transient and stable cell clusters every 30 min for 5.5 h post inoculation in 12 random fields of view for each strain. We were unable to follow the fate of cell clusters beyond this time point in this assay due to the high density of planktonic cells that obscured the cell clusters. A cell cluster was defined as being stable if it was observed in at least two consecutive time points and also still present at 5.5 h. Transient cell clusters were defined as being present in at least one time point but were absent at the final time-point (5.5 h).

We have shown previously that eDNA released during the initial stages of submerged biofilm formation occurs via Lys-mediated explosive cell lysis (13). Here we found that the majority of these eDNA release events rapidly dissipated and were not associated with the formation of cell clusters visible within the time-frames captured (Figure 3). However, by the end of the time period (5.5 h) across all 12 fields of view, PAO1 formed a total of 24 stable and 2 transient cell clusters, PAO1Δ*alpB* formed 9 stable and 4 transient cell clusters, PAO1Δ*cidAB* formed no stable cell clusters and 2 transient cell clusters and PAO1Δ*hol* formed 8 stable and 10 transient cell clusters (Figure 3; Figure S5-S6).

**Figure 3.**
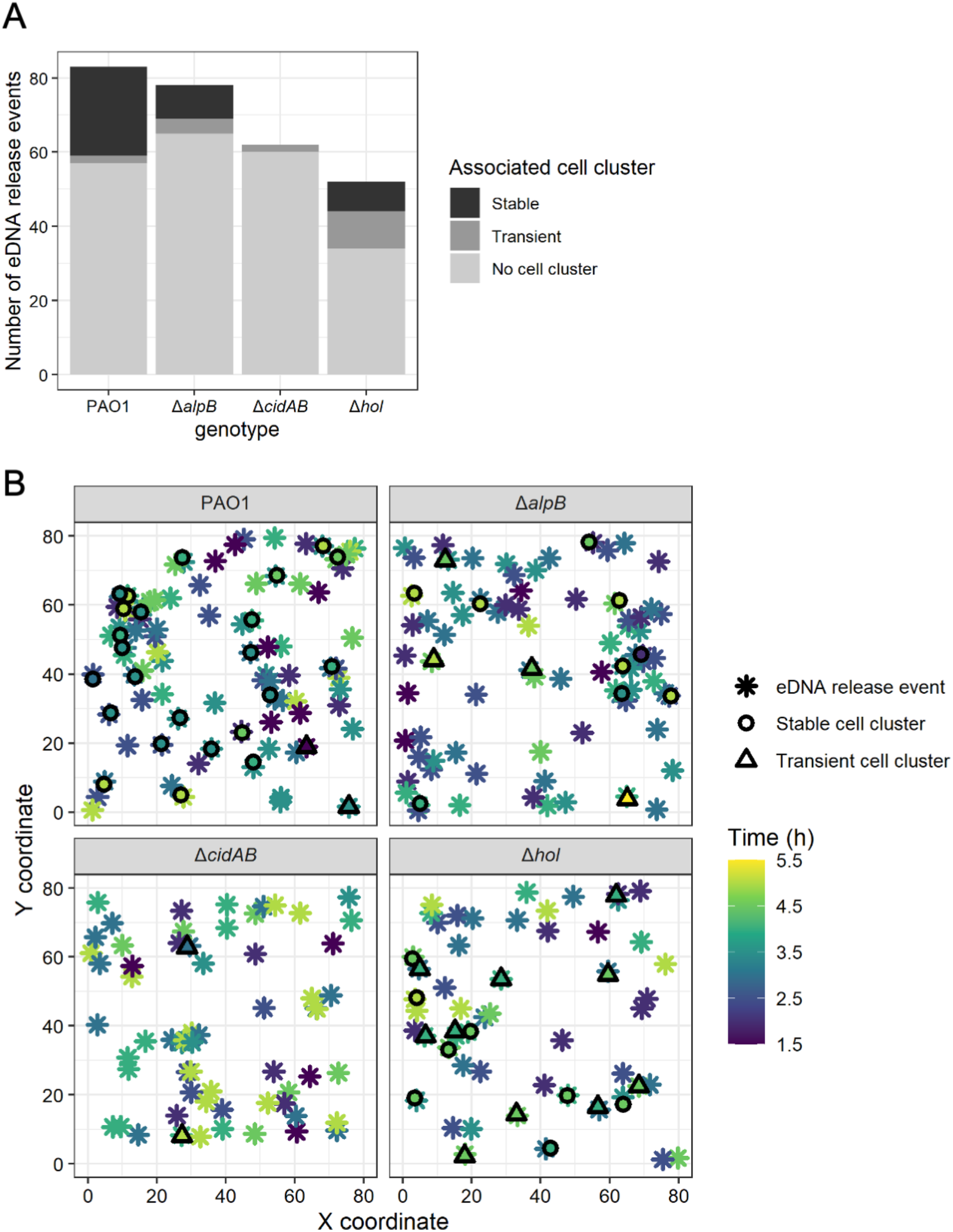
eDNA release events initiate the formation of cell clusters in submerged biofilms. Cell clusters and eDNA (as visualised with TOTO-1 stain) release events by PAO1, PAO1Δ*alpB,* PAO1Δ*cidAB* and PAO1Δ*hol* from 1-5.5 h at 37°C. (A) Total eDNA release events that are associated with stable cell clusters, transient cell clusters or not associated with any cell clusters. (B) X, Y coordinates for eDNA release events and associated stable or transient cell clusters. Data are combined from 12 random fields of view across 2 biological replicates (n=12). See Figures S5-6 for data by biological replicate and field of view.

We also explored the temporal relationship between eDNA release and the formation of stable and transient cell clusters. For all strains, the sites of stable or transient cell clustering were associated with an eDNA release event either within the same 30 min period or prior to the cluster formation (Figure 3B; Figure 4A; Figure S5-S6) with the majority of stable cell clusters forming within 1 h of a prior eDNA release event at the same location (Figure 3B; Figure S5-S6). These observations suggest that eDNA released via explosive cell lysis initiates subsequent cell clustering and microcolony formation during the early stages of submerged biofilm development.

**Figure 4.**
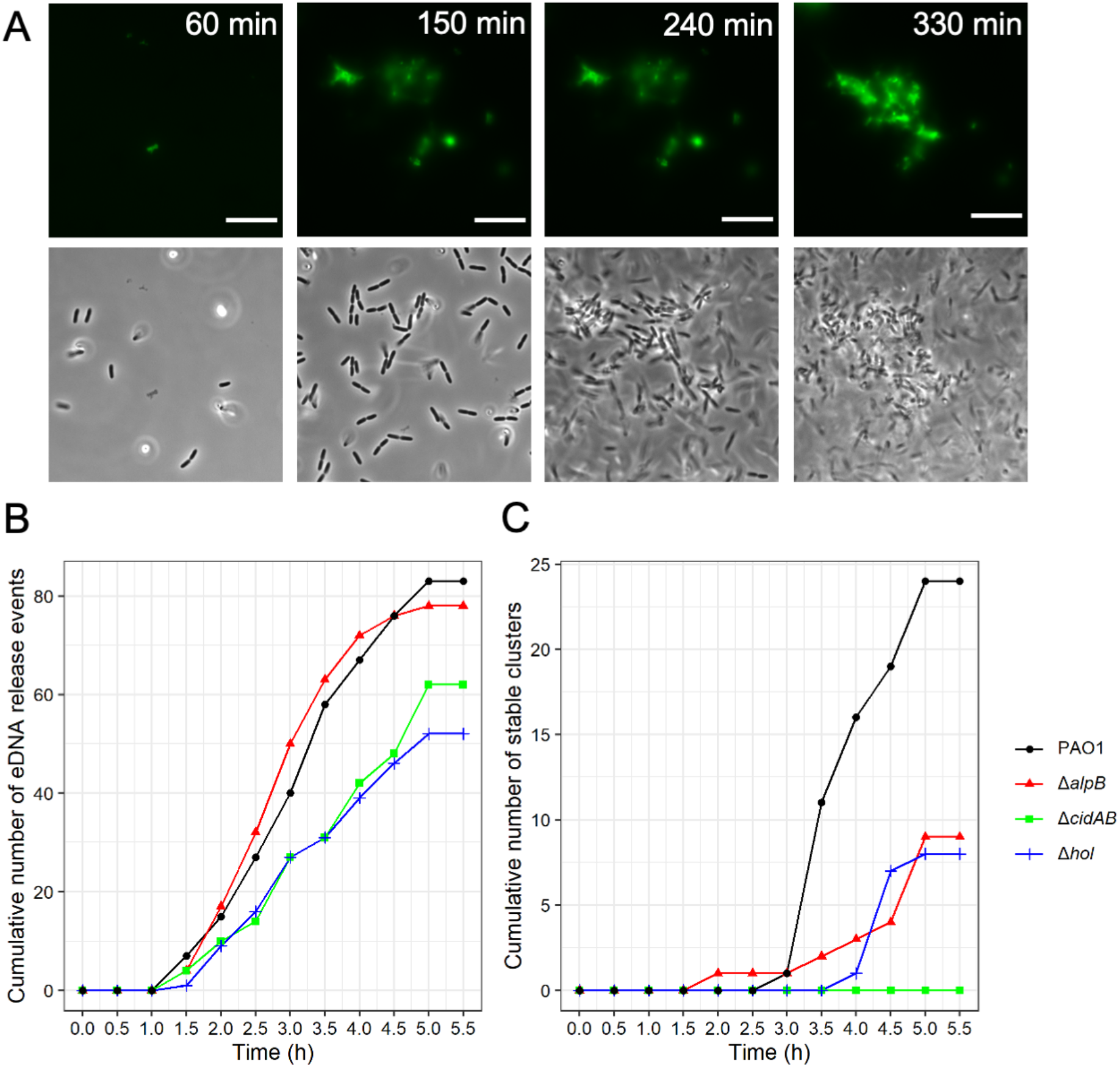
The ability to release eDNA is not sufficient for stable cell clusters to form. (A) eDNA release events and associated stable cell cluster formation by PAO1 with phase contrast (bottom) and eDNA (TOTO-1, top); Scale bar=10 μm. eDNA release events from 1-5.5 hr for PAO1, PAO1Δ*alpB,* PAO1Δ*cidAB* and PAO1Δ*hol* represented as (B) cumulative sum of all eDNA release events or (C) the cumulative sum of stable cell clusters from 1-5.5 h. Data are combined from 12 random fields of view across 2 biological replicates (n=12). Means and 95% CIs were estimated using a negative binomial regression of event counts at 5.5 h on genotype. See Table S8-9 for associated effect sizes, 95% CIs and p-values.

To determine why the holin mutants were either completely (PAO1Δ*cidAB*) or partially (PAO1Δ*alpB* and PAO1Δ*hol*) defective in the formation of stable cell clusters we looked at the total number of eDNA release events, including those that were not associated with cell clusters, and when these events occurred. In all strains, including PAO1, the first eDNA events occurred 1.5 h after inoculation (Figure 4B). Over the ensuing 4.5 h PAO1 had a total of 83 eDNA release events at distinct locations. There was no significant difference in the total number of eDNA release events for PAO1Δ*alpB* and PAO1Δ*cidAB* compared to PAO1 (78 events, p=0.77 and 62 events, p=0.162 for PAO1Δ*alpB* and PAO1Δ*cidAB* respectively, compared to PAO1; Table S8). Only PAO1Δ*hol* showed significantly fewer total eDNA events (52 events, p=0.039; Table S8) compared to PAO1 (Figure 4B).

These observations suggest that the ability to release eDNA is not sufficient for stable cell clusters to form, as PAO1Δ*cidAB* is able to release eDNA at a similar rate as PAO1 but did not form any stable cell clusters during this 5.5 h period (Figure 4B). Additionally, while both PAO1Δ*alpB* and PAO1Δ*hol* were also able to release eDNA, their ability to form stable cell clusters was significantly impaired compared to PAO1 (rate ratios=0.37 or 0.33; 95% CI=0.17-0.81 or 0.15-0.74; p=0.012 or 0.007 for PAO1Δ*alpB* and PAO1Δ*hol*, respectively; Table S9), with both of these mutants forming stable cell clusters at approximately 1/3 the rate of PAO1 (Figure 4C). While PAO1Δ*cidAB* did not form any stable cell clusters during the first 5.5 h this strain did form some microcolonies by 8 h (Figure 2), which suggests that in the period between 5.5 h to 8 h in the absence of CidA the two other holins, AlpB and Hol, are eventually able to contribute sufficient eDNA to form stable cell clusters and develop microcolonies.

### Cell cluster consolidation requires a continual increase in eDNA levels over time

One consideration that would affect the ability to consolidate a stable cell cluster and influence the size of the subsequent microcolony, is how much eDNA accumulates within the cell cluster over time. Indeed, at 8 h the amount of eDNA in the microcolony appears to correlate with the size of the microcolony (Figure S3). To investigate this further, we followed the accumulation of eDNA in each stable cell cluster in the 24 PAO1, 9 PAO1Δ*alpB* and 8 PAO1Δ*hol* stable cell clusters by evaluating the integrated density value of the corresponding fluorescence signal of the eDNA stained with TOTO-1 over the time period of 1-5.5 h. We then modelled the growth of integrated density values over time since the first appearance of a cluster as a function of genotype (Figure 5A). These analyses showed that there was significant variation in trajectories between clusters according to genotype. While PAO1 and PAO1Δ*alpB* clusters showed similar increases in integrated densities over time, there was significantly less increase in integrated densities for PAO1Δ*hol* over time, and in fact an overall decrease in many cases (Figure 5A; Table S10). Together these data suggest that both PAO1 and PAO1Δ*alpB* are able to provide sustained release of eDNA over time within the cell cluster, whereas PAO1Δ*hol* is partially defective in this process.

**Figure 5.**
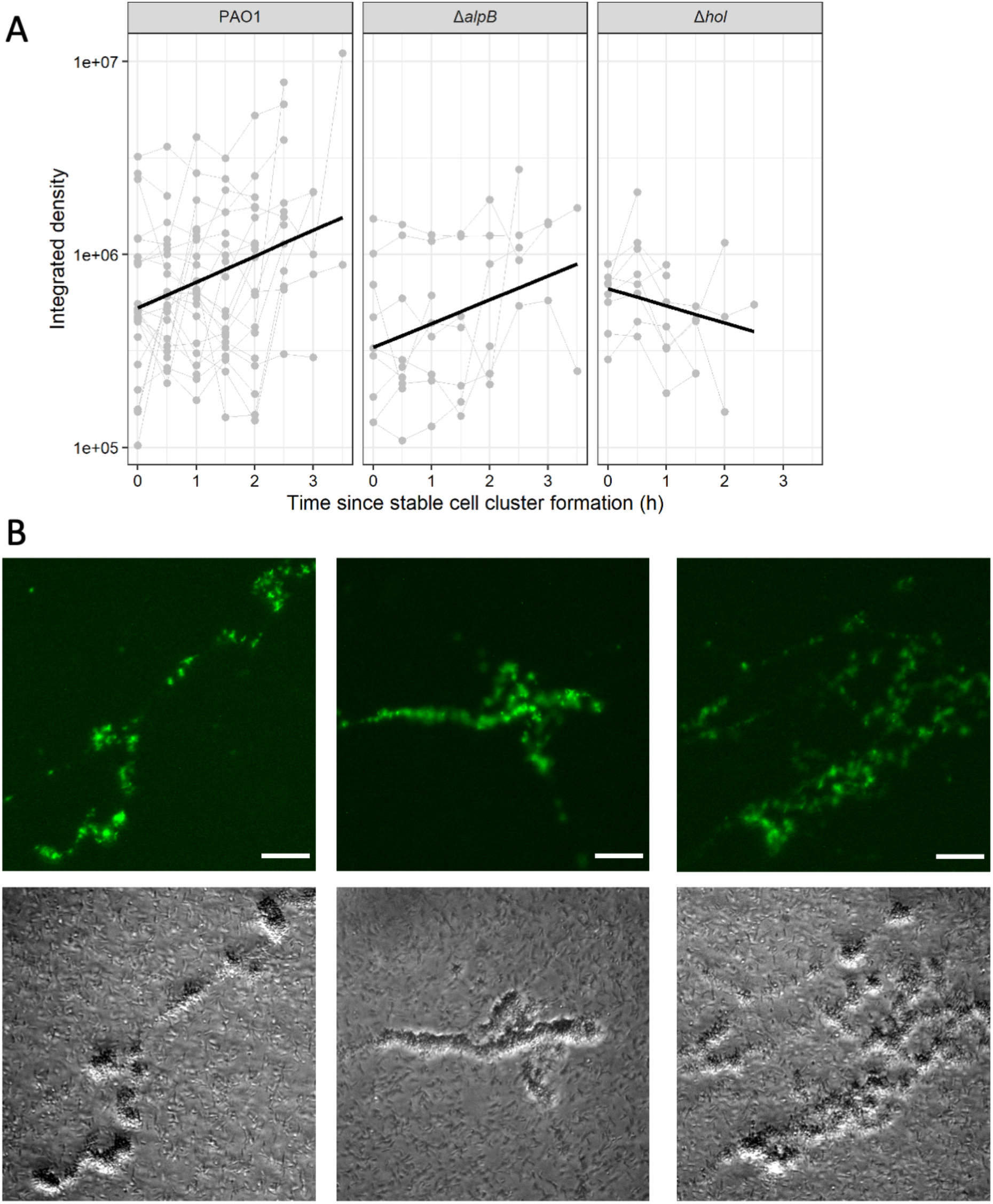
Holins are involved in sustained eDNA release over time to consolidate stable cell clusters. (A) Integrated density of fluorescence signal (eDNA – as visualised with TOTO-1 stain) associated with stable cell clusters formed over 1-5.5 h for PAO1, PAO1Δ*alpB* or PAO1Δ*hol.* Original data was generated from analysis of 12 random fields of view across 2 biological replicates (n=12). Slopes correspond to average growth over time, estimated using a linear mixed model with log(density) as an outcome, time, genotype and their interaction as fixed effects, and random slope of density over time. The slope of PAO1Δ*hol* is significantly different from PAO1 - see Table S10 for log growth rate, 95% CIs and p-values. (B) Multiple eDNA release events within the same or adjacent stable cell clusters results in the formation of large multi-structured microcolonies in PAO1 submerged biofilms after incubation at 37°C for 8 h. Images are eDNA (EthHD-2, top) or phase contrast (bottom). Scale=10 μm.

We noted that in 4 of the 24 PAO1 stable cell clusters, the amount of eDNA accumulated to a much higher level than in other clusters (Figure 5A). For these cell clusters it appeared that in addition to the initial explosive lysis event, at least one additional large release event occurred within the cell cluster. It is likely that multiple explosive events within the same cell cluster and/or within adjacent cell clusters may explain how many very large microcolony structures were observed at 8 h in PAO1 (69/203 (33%) microcolonies were greater than 200 μm^2^) (Figure 5B; Figure 2C) but at a much lower frequency in PAO1Δ*alpB,* PAO1Δ*cidAB* or PAO1Δ*hol* (5/65 (8%), 1/42 (2%) or 2/107 (2%) were greater than 200 μm^2^) (Figure 2C).

We also followed the integrated density values of TOTO-1 stained eDNA fluorescence over time for transient cell clusters of each strain and observed that in every case the eDNA content of the transient clusters had dissipated within 30 min of the initial eDNA release event (Figure S7). Overall our observations suggest that cell cluster consolidation requires sustained release of eDNA over the early stages of biofilm formation and that releasing more eDNA is likely to result in larger microcolonies.

## Discussion

In this study we found that eDNA release through explosive cell lysis determines the sites at which cells begin to cluster to form microcolonies in later stages of submerged biofilm development. The spatial overlap between sites of eDNA release and sites of cell clusters was 100% and it appeared that cell clusters never formed in the absence of a prior eDNA release event. Our observations suggest that eDNA released through Lys-mediated explosive cell lysis is required for the initiation of microcolony formation in submerged biofilms.

We investigated the contribution of the putative *P. aeruginosa* holins Hol, AlpB and CidA to eDNA release within *P. aeruginosa* interstitial and submerged biofilms. We found that while all 3 holins are able to facilitate Lys-mediated eDNA release, their contribution to this varies depending upon the type of biofilm and the stage of biofilm development. We found that AlpB and Hol appear to be the key holins for eDNA release in actively expanding interstitial biofilms. In contrast, within submerged biofilms, all 3 holins are involved in eDNA release during biofilm development to different extents. Specifically, PAO1Δ*hol* was the only holin mutant significantly defective in explosive cell lysis events during the initial 5.5 h period (Figure 3, Figure 4B-C, Table S8). This strain also had more transient cell clusters and significantly fewer stable cell clusters than PAO1 and appeared to trend toward losing eDNA in the stable cell clusters over time (Figure 3, Figure 4B-C, Figure 5A). These observations suggest that Hol is involved in the initial release of eDNA via explosive cell lysis and also contributes to the sustained release of eDNA during cluster consolidation. In contrast, in the absence of CidA there was no significant defect in the rate of eDNA release events during the first 5.5 h (Figure 3, Figure 4B, Table S8). However, PAO1Δ*cidAB* failed to form any stable cell clusters during this period (Figure 3). This suggests that CidA is not involved in the initial release of eDNA via explosive cell lysis but does contribute to the sustained release of eDNA that is required to stabilise clusters during the initial stages of submerged biofilm formation. Finally, PAO1Δ*alpB* did not show significant defects in the rate of eDNA release events but was significantly defective in the formation of stable cell clusters (Figure 3, Figure 4B-C, Table S9). However, unlike PAO1Δ*hol,* the stable cell clusters formed by PAO1Δ*alpB* tended to increase eDNA content in the stable cell clusters as measured by integrated fluorescence intensity (Figure 5A). Despite this increase in eDNA content in stable clusters, the microcolonies formed by PAO1Δ*alpB* at 8 h were significantly smaller than those of PAO1 (Figure 2C, Table S5), which suggests that AlpB contributes to further release of eDNA after stable cell cluster formation when cells begin to form microcolonies. Due to the defects in stable cluster formation by PAO1Δ*hol* and PAO1Δ*cidAB* we are unable to determine the contribution of Hol and AlpB to microcolony development following stable cluster formation.

One hypothesis to explain our observations is that each holin is responsible for eDNA release at a different time during submerged biofilm development. This is supported by the known role for holins as the ‘protein clocks’ of bacteriophage infections (34). Although lysis is executed by the action of a muralytic endolysin, in this case Lys, it is temporally controlled by a cognate holin (23). Lysis occurs when holin expression is triggered, at a specific time that is “programmed” into the holin gene, to form endolysin-transporting pores in the inner membrane (34). It is possible that each of AlpB, CidA and Hol have different “programmed” times of activity during biofilm development and as such contribute differing roles during biofilm development. It may also be that each holin is expressed at different levels in different stages of submerged biofilm development.

It is also possible that different environmental signals determine how much of each holin will be expressed and when this will occur. This may account for why different holins are involved in eDNA release in actively expanding or submerged biofilms and at different stages of submerged biofilm development. We do not currently know what controls holin and Lys-mediated explosive cell lysis in *P. aeruginosa.* While we have previously demonstrated that exogenous stress increases the frequency of cells undergoing explosive cell lysis, exogenous stress *per se* is not required for this phenomenon to occur (19). Previous studies have shown both *hol* and *lys* are under the regulation of RecA and PrtN (20), *alpB* expression to not be affected by deletion of *prtN* and instead to be regulated by AlpR (24) and *cidA* to be activated by CidR, which is upstream of CidA (11). It is possible that each of these holins is regulated by these proteins to mediate explosive cell lysis in response to certain conditions however, it is also possible that there is some overlap or crosstalk between these pathways, or even that regulation of holins for explosive cell lysis is controlled in a completely different manner.

Given that explosive cell lysis releases all cellular contents as public goods (19) a temporally regulated, partially redundant holin system for explosive cell lysis would support the controlled production and distribution of other essential biofilm matrix components. Our data in this study demonstrates that sustained eDNA release over time is required for cell cluster consolidation and subsequent microcolony formation. This may fit with the ‘rich get richer’ model that Zhao *et al* proposed (35) for another matrix component, the exopolysaccharide Psl. In their study they propose that Psl is laid down by single cells and as cells cluster around the Psl, they produce more Psl, creating a positive feedback loop. It has been shown that Psl and eDNA physically interact in established submerged biofilms (36). It would therefore be interesting to determine in future work where Psl trails and eDNA are relative to one another in the very early stages of submerged biofilm development and also if eDNA release by explosive cell lysis attracts more cells to the cell cluster, which then leads to more eDNA release, facilitating microcolony formation.

One additional consideration for why each of AlpB, CidA and Hol have different roles in actively expanding and submerged biofilms, and why they are involved in different stages of submerged biofilm formation may also be linked to putative interactions with spanins, another component of the phage lysis system. These proteins accumulate in the inner and outer membranes and are involved in rupturing the outer membrane of Gram negative bacteria to complete the lysis process (23). However, little work has been done on these proteins in *P. aeruginosa* and it is unknown whether holins and spannins interact in the inner membrane, when they are expressed, or if spanins play any role in explosive cell lysis. Indeed, investigation of these possibilities would be an interesting avenue to explore in future work.

The available options for treating bacterial biofilms are becoming increasingly limited. Fully understanding the spatial, temporal and mechanical control of biofilm formation, as well as the genetic components involved, will assist with the development of novel strategies to prevent and disrupt biofilms. In the current study we show that multiple holins contribute to Lys-mediated explosive cell lysis in both interstitial and submerged *P. aeruginosa* biofilms. We also demonstrate that eDNA release occurs prior to microcolony initiation and that sustained eDNA release is required for microcolony development. This adds to our understanding of how *P. aeruginosa* biofilm development occurs and could aid in the development of antimicrobial treatments targeting this process.

## Authors and Contributions

A.L.H and J.J.L conducted experiments. A.L.H, G.M.S, L.M.N and C.B.W analysed results. L.T, L.M.N and C.B.W provided project supervision. C.B.W provided project administration and funding. A.L.H, L.C.M, L.M.N and C.B.W wrote the manuscript.

## References

1. Donlan RM, Costerton JW. Biofilms: survival mechanisms of clinically relevant microorganisms. Clin Microbiol Rev. 2002 Apr;15(2):167–93.

2. Donlan RM. Biofilms and device-associated infections. Emerging Infect Dis. 2001 Apr;7(2):277–81.

3. Hughes G, Webber MA. Novel approaches to the treatment of bacterial biofilm infections. Br J Pharmacol. 2017 Jul;174(14):2237–46.

4. Xu D, Jia R, Li Y, Gu T. Advances in the treatment of problematic industrial biofilms. World J Microbiol Biotechnol. 2017 May;33(5):97.

5. Flemming H-C, Wingender J. The biofilm matrix. Nat Rev Microbiol. 2010 Sep;8(9):623–33.

6. Payne DE, Boles BR. Emerging interactions between matrix components during biofilm development. Curr Genet. 2016 Feb;62(1):137–41.

7. Vu B, Chen M, Crawford RJ, Ivanova EP. Bacterial extracellular polysaccharides involved in biofilm formation. Molecules. 2009 Jul 13;14(7):2535–54.

8. Okshevsky M, Meyer RL. The role of extracellular DNA in the establishment, maintenance and perpetuation of bacterial biofilms. Crit Rev Microbiol. 2013 Dec 4;41(3):341–52.

9. Southey-Pillig CJ, Davies DG, Sauer K. Characterization of temporal protein production in *Pseudomonas aeruginosa* biofilms. J Bacteriol. 2005 Dec;187(23):8114–26.

10. Palmer J, Flint S, Brooks J. Bacterial cell attachment, the beginning of a biofilm. J Ind Microbiol Biotechnol. 2007 Sep;34(9):577–88.

11. Ma L, Conover M, Lu H, Parsek MR, Bayles K, Wozniak DJ. Assembly and development of the *Pseudomonas aeruginosa* biofilm matrix. PLoS Pathog. 2009 Mar 27;5(3):e1000354.

12. McDougald D, Klebensberger J, Tolker-Nielsen T, Webb JS, Conibear T, Rice SA, et al. Pseudomonas aeruginosa: A Model for Biofilm Formation. In: Rehm BHA, editor. Pseudomonas. Weinheim, Germany: Wiley-VCH Verlag GmbH & Co. KGaA; 2008. p. 215–53.

13. Whitchurch CB, Tolker-Nielsen T, Ragas PC, Mattick JS. Extracellular DNA required for bacterial biofilm formation. Science. 2002 Feb 22;295(5559):1487.

14. Gloag ES, Turnbull L, Huang A, Vallotton P, Wang H, Nolan LM, et al. Self-organization of bacterial biofilms is facilitated by extracellular DNA. Proc Natl Acad Sci USA. 2013 Jul 9;110(28):11541–6.

15. Fuxman Bass JI, Russo DM, Gabelloni ML, Geffner JR, Giordano M, Catalano M, et al. Extracellular DNA: a major proinflammatory component of *Pseudomonas aeruginosa* biofilms. J Immunol. 2010 Jun 1;184(11):6386–95.

16. Mulcahy H, Charron-Mazenod L, Lewenza S. Extracellular DNA chelates cations and induces antibiotic resistance in *Pseudomonas aeruginosa* biofilms. PLoS Pathog. 2008 Nov 21;4(11):e1000213.

17. Madsen JS, Burmølle M, Hansen LH, Sørensen SJ. The interconnection between biofilm formation and horizontal gene transfer. FEMS Immunol Med Microbiol. 2012 Jul;65(2):183–95.

18. Ibáñez de Aldecoa AL, Zafra O, González-Pastor JE. Mechanisms and regulation of extracellular DNA release and its biological roles in microbial communities. Front Microbiol. 2017 Jul 26;8:1390.

19. Turnbull L, Toyofuku M, Hynen AL, Kurosawa M, Pessi G, Petty NK, et al. Explosive cell lysis as a mechanism for the biogenesis of bacterial membrane vesicles and biofilms. Nat Commun. 2016 Apr 14;7:11220.

20. Nakayama K, Takashima K, Ishihara H, Shinomiya T, Kageyama M, Kanaya S, et al. The R-type pyocin of *Pseudomonas aeruginosa* is related to P2 phage, and the F-type is related to lambda phage. Mol Microbiol. 2000 Oct;38(2):213–31.

21. Amini S, Hottes AK, Smith LE, Tavazoie S. Fitness landscape of antibiotic tolerance in *Pseudomonas aeruginosa* biofilms. PLoS Pathog. 2011 Oct 20;7(10):e1002298.

22. Saier MH, Reddy BL. Holins in bacteria, eukaryotes, and archaea: multifunctional xenologues with potential biotechnological and biomedical applications. J Bacteriol. 2015 Jan 1;197(1):7–17.

23. Young R. Phage lysis: three steps, three choices, one outcome. J Microbiol. 2014 Mar 1;52(3):243–58.

24. McFarland KA, Dolben EL, LeRoux M, Kambara TK, Ramsey KM, Kirkpatrick RL, et al. A self-lysis pathway that enhances the virulence of a pathogenic bacterium. Proc Natl Acad Sci USA. 2015 Jul 7;112(27):8433–8.

25. Bayles KW. Are the molecular strategies that control apoptosis conserved in bacteria? Trends Microbiol. 2003 Jul;11(7):306–11.

26. Rice KC, Mann EE, Endres JL, Weiss EC, Cassat JE, Smeltzer MS, et al. The *cidA* murein hydrolase regulator contributes to DNA release and biofilm development in *Staphylococcus aureus*. Proc Natl Acad Sci USA. 2007 May 8;104(19):8113–8.

27. Ranjit DK, Endres JL, Bayles KW. *Staphylococcus aureus* CidA and LrgA proteins exhibit holin-like properties. J Bacteriol. 2011 May;193(10):2468–76.

28. Hoang TT, Karkhoff-Schweizer RR, Kutchma AJ, Schweizer HP. A broad-host-range Flp-FRT recombination system for site-specific excision of chromosomally-located DNA sequences: application for isolation of unmarked *Pseudomonas aeruginosa* mutants. Gene. 1998 May 28;212(1):77–86.

29. Toyofuku M, Zhou S, Sawada I, Takaya N, Uchiyama H, Nomura N. Membrane vesicle formation is associated with pyocin production under denitrifying conditions in *Pseudomonas aeruginosa* PAO1. Environ Microbiol. 2014 Sep;16(9):2927–38.

30. Schindelin J, Arganda-Carreras I, Frise E, Kaynig V, Longair M, Pietzsch T, et al. Fiji: an open-source platform for biological-image analysis. Nat Methods. 2012 Jun 28;9(7):676–82.

31. Bates D, Mächler M, Bolker B, Walker S. Fitting Linear Mixed-Effects Models Using lme4. J Stat Softw. 2015;67(1).

32. Kuznetsova A, Brockhoff PB, Christensen RHB. lmertest package: tests in linear mixed effects models. J Stat Softw. 2017;82(13).

33. https://www.R-project.org/ RCT (2019). R: A language and environment for statistical computing. R Foundation for Statistical Computing, Vienna, Austria; 2019.

34. Wang IN, Smith DL, Young R. Holins: the protein clocks of bacteriophage infections. Annu Rev Microbiol. 2000;54:799–825.

35. Zhao K, Tseng BS, Beckerman B, Jin F, Gibiansky ML, Harrison JJ, et al. Psl trails guide exploration and microcolony formation in Pseudomonas aeruginosa biofilms. Nature. 2013 May 16;497(7449):388–91.

36. Wang S, Liu X, Liu H, Zhang L, Guo Y, Yu S, et al. The exopolysaccharide Psl-eDNA interaction enables the formation of a biofilm skeleton in Pseudomonas aeruginosa. Environ Microbiol Rep. 2015 Apr;7(2):330–40.

