## Supplementary Tables and Figures for "Multiple Holins Contribute to Extracellular DNA Release in *Pseudomonas aeruginosa* Biofilms"

† These authors contributed equally

### Supplementary Tables S1-S10 and Supplementary Figures S1-S7

### Supplementary Tables

**Table S1. Strains and plasmids used in this study**

| Strains | Relevant characteristics | Source |
|---|---|---|
| ***E. coli*** | | |
| DH5α | *recA, endA1, gyrA96, hsdR17, thi-1, supE44, relA1*, φ80, dlacZΔM15 | Laboratory collection |
| S17-1 | Donor strain for conjugal transfer (*thi pro hsdR recA chr::RP4-2*) | (1) |
| ***P. aeruginosa*** | | |
| PAO1 | Wild type *P. aeruginosa* strain | (2) |
| PAO1Δ*alpB* | In frame deletion of *alpB* (PA0908) | This study |
| PAO1Δ*cidAB* | In frame deletion of *cidAB* (PA3432-3431) | (2) |
| PAO1Δ*hol* | In frame deletion of *hol* (PA0614) | This study |
| PAO1Δ*cidAB*Δ*alpB* | In frame deletion of *cidAB* and *alpB*. Parent strain PAO1Δ*cidAB* | This study |
| PAO1Δ*cidAB*Δ*hol* | In frame deletion of *cidAB* and *hol.* Parent strain PAO1Δ*cidAB* | This study |
| PAO1Δ*hol*Δ*alpB* | In frame deletion of *hol* and *alpB*. Parent strain PAO1Δ*hol* | This study |
| PAO1Δ*cidAB*Δ*hol*Δ*alpB* | In frame deletion of *cidAB*, *hol* and *alpB.* Parent strain PAO1Δ*cidAB*Δ*hol* | This study |
| PAO1Δ*lys* | In frame deletion of *lys* | (3) |
| **Plasmids** | **Relevant characteristics** | **Source** |
| pJN105 | Broad host range arabinose inducible gene expression vector ($Gm^R$) | (4) |
| pJN*alpB* | pJN105 with wild type *alpB* ($Gm^R$) | This study |
| pJN*cidAB* | pJN105 with wild type *cidAB* ($Gm^R$) | This study |
| pJN*hol* | pJN105 with wild type *hol* ($Gm^R$) | This Study |
| pJN*lys* | pJN105 with wild type *lys* ($Gm^R$) | (3) |
| pRIC380 | Suicide vector ($Amp^R$) | (5) |
| pPS856 | *FRT* cassette vector ($Gm^R$) | (6) |
| pFLP2 | Site-specific excision vector ($Gm^R$) | (6) |
| pALH3 | 1kb upstream and 1kb downstream flanking regions of *hol* with a *Hind*III site in between and flanking *Spe*I sites, synthesised in pMK-RQ; $Km^R$ | This study |
| pALH5 | pALH3 containing *FRT-Gm$^R$-FRT* from pPS856 cloned into *Hind*III site; $Gm^R$, $Km^R$ | This study |
| pALH7 | 3.1 kb *Spe*I insert from pALH5 cloned into pRIC380; $Gm^R$, $Ap^R$ | This study |
| pALH9 | 1kb upstream and 1kb downstream flanking regions of *alpB* with a *Hind*III site in between and flanking *Spe*I sites, synthesised in pMK-RQ; $Km^R$ | This study |
| pALH10 | pALH9 containing *FRT-Gm$^R$-FRT* from pPS856 cloned into *Hind*III site; $Gm^R$, $Km^R$ | This study |
| pALH11 | 3.1 kb *Spe*I insert from pALH10 cloned into pRIC380; $Gm^R$, $Ap^R$ | This study |

$Gm^R$ - Gentamicin resistance, $Km^R$ - Kanamycin resistance, $Ap^R$ - Ampicillin resistance

**Table S2. Primers used in this study**

| Primers | Sequence (5' to 3') |
|---|---|
| alpB_F | TGACGCTATGGGACGATAAA |
| alpB_R | GTACGTTCGTTCAATGCAGG |
| cidAB_F | CTTTCTCCATCCCCGATTTC |
| cidAB_R | TTTTGTCGTTATCGGATGCC |
| hol_F | TTCTTGTAAAGGTGCGTCCC |
| hol_R | GCATGGTTGACTCCTTCGAT |

**Table S3.** Incidence rate ratios for the number of eDNA release events in interstitial biofilms per unit area, with 95% confidence intervals and p-values (corresponding to the data shown in Figure 1C) with each mutant compared to PAO1.

| Genotype | Incidence Rate Ratio* | 95% CI | p-value |
|---|---|---|---|
| Δ*alpB* | 0.9 | 0.52 – 1.55 | 0.702 |
| Δ*cidAB* | 0.9 | 0.52 – 1.55 | 0.705 |
| Δ*hol* | 1.08 | 0.63 – 1.86 | 0.768 |
| Δ*alpB*Δ*cidAB* | 1.02 | 0.60 – 1.76 | 0.929 |
| Δ*alpB*Δ*hol* | 0.06 | 0.03 – 0.10 | **<0.001** |
| Δ*cidAB*Δ*hol* | 0.85 | 0.49 – 1.46 | 0.556 |
| Δ*hol*Δ*cidAB*Δ*alpB* | 0.07 | 0.04 – 0. 12 | **<0.001** |
| Δ*lys* | 0.04 | 0.02 – 0.08 | **<0.001** |

* rate of eDNA events/area analysed, compared to PAO1

**Table S4.** Incidence rate ratios of average number of microcolonies in submerged biofilms between genotypes with 95% confidence intervals and p-values (corresponding to the data shown in Figure 2B).

| Genotype comparison | Incidence Rate Ratio* | 95% CI | p-value |
|---|---|---|---|
| PAO1 – Δ*alpB* | 3.86 | 2.86 - 5.2 | **<0.001** |
| PAO1 – Δ*cidAB* | 5.80 | 4.09 - 8.23 | **<0.001** |
| PAO1 – Δ*hol* | 2.35 | 1.82 - 3.04 | **<0.001** |
| Δ*alpB* – Δ*cidAB* | 1.50 | 1.01 - 2.25 | 0.047 |
| Δ*alpB* – Δ*hol* | 0.61 | 0.44 - 0.842 | 0.003 |
| Δ*cidAB* – Δ*hol* | 0.40 | 0.279 - 0.587 | **<0.001** |

* number of microcolonies of first strain listed/second strain listed

**Table S5.** Incidence rate ratios of average microcolony size between genotypes with 95% confidence intervals and p-values (corresponding to the data shown in Figure 2C).

| Genotype comparison | Incidence Rate Ratio* | 95% CI | p-value |
|---|---|---|---|
| PAO1 – Δ*alpB* | 1.84 | 1.39 - 2.45 | **<0.001** |
| PAO1 – Δ*cidAB* | 2.71 | 1.94 - 3.78 | **<0.001** |
| PAO1 – Δ*hol* | 2.33 | 1.82 - 2.99 | **<0.001** |
| Δ*alpB* – Δ*cidAB* | 1.47 | 1.01 - 2.15 | 0.047 |
| Δ*alpB* – Δ*hol* | 1.26 | 0.93 - 1.72 | 0.133 |
| Δ*cidAB* – Δ*hol* | 0.86 | 0.605 - 1.22 | 0.401 |

* microcolony size of first strain listed/second strain listed

**Table S6.** Incidence rate ratios of average integrated density of TOTO-1 fluorescence in 8 h microcolonies between genotypes with 95% confidence intervals and p-values (corresponding to the data shown in Figure 2D).

| Genotype comparison | Incidence Rate Ratio* | 95% CI | p-value |
|---|---|---|---|
| PAO1 – Δ*alpB* | 3.48 | 2.03 - 5.95 | **<0.001** |
| PAO1 – Δ*cidAB* | 4.43 | 2.38 - 8.24 | **<0.001** |
| PAO1 – Δ*hol* | 6.97 | 4.29 - 11.3 | **<0.001** |
| Δ*alpB* – Δ*cidAB* | 1.27 | 0.643 - 2.52 | 0.485 |
| Δ*alpB* – Δ*hol* | 2.01 | 1.14 - 3.53 | **0.016** |
| Δ*cidAB* – Δ*hol* | 1.57 | 0.828 - 2.99 | 0.165 |

* integrated density of first strain listed/second strain listed

**Table S7.** Incidence rate ratios of average number of microcolonies between genotypes with 95% confidence intervals and p-values (corresponding to the data shown in Figure S4B).

| Genotype comparison | Incidence Rate Ratio* | 95% CI | p-value |
|---|---|---|---|
| PAO1 [pJN105] – Δ*alpB* [pJN105] | 0.58 | 0.33 – 1.01 | 0.055 |
| PAO1 [pJN105] – Δ*cidAB* [pJN105] | 0.39 | 0.21 – 0.75 | **0.004** |
| PAO1 [pJN105] – Δ*hol* [pJN105] | 0.48 | 0.27 – 0.88 | **0.017** |
| PAO1 [pJN105] – PAO1 [pJN*alpB*] | 0.79 | 0.47 – 1.32 | 0.363 |
| PAO1 [pJN105] – PAO1 [pJN*cidAB*] | 0.76 | 0.45 – 1.27 | 0.295 |
| PAO1 [pJN105] – PAO1 [pJN*hol*] | 0.73 | 0.43 – 1.23 | 0.235 |
| Δ*alpB* [pJN105] – Δ*alpB* [pJN*alpB*] | 2.74 | 1.30 – 5.79 | **0.008** |
| Δ*cidAB* [pJN105] – Δ*cidAB* [pJN*cidAB*] | 2.44 | 1.04 – 5.71 | **0.04** |
| Δ*hol* [pJN105] – Δ*hol* [pJN*hol*] | 2.84 | 1.28 – 6.28 | **0.01** |

* number of microcolonies of first strain listed/second strain listed

**Table S8.** Incidence rate ratios for the rate of all eDNA release events over time, with 95% confidence intervals and p-values (corresponding to the data shown in Figure 4B) with each mutant compared to PAO1.

| Genotype | Incidence Rate Ratio* | 95% CI | p-value |
|---|---|---|---|
| Δ*alpB* | 0.94 | 0.62 – 1.42 | 0.77 |
| Δ*cidAB* | 0.73 | 0.48 – 1.13 | 0.162 |
| Δ*hol* | 0.63 | 0.40 – 0.98 | **0.039** |

* eDNA events for mutant/eDNA events for PAO1

**Table S9.** Incidence rate ratios for the rate of eDNA release events associated with stable cell clusters over time, with 95% confidence intervals and p-values (corresponding to the data shown in Figure 4C) with each mutant compared to PAO1.

| Genotype | Incidence Rate Ratio* | 95% CI | p-value |
|---|---|---|---|
| Δ*alpB* | 0.37 | 0.17 – 0.81 | **0.012** |
| Δ*cidAB* | 0 | - | - |
| Δ*hol* | 0.33 | 0.15 – 0.74 | **0.007** |

* eDNA events for mutant/eDNA events for PAO1

**Table S10.** Log growth rate for integrated density of eDNA (fluorescent signal) in stable cell clusters over time with t=0 being when the cluster first appeared. The 95% confidence intervals and p-values (corresponding to the data shown in Figure 5A) with each mutant compared to PAO1.

| Genotype | Log growth rate | 95% CI | p-value |
|---|---|---|---|
| PAO1 | 0.31 | 0.13 – 0.49 | **-** |
| Δ*alpB* | 0.29 | 0.00 – 0.58 | **0.9896** |
| Δ*hol* | -0.21 | -0.58 – 0.17 | **0.0432** |

### Supplementary Figures

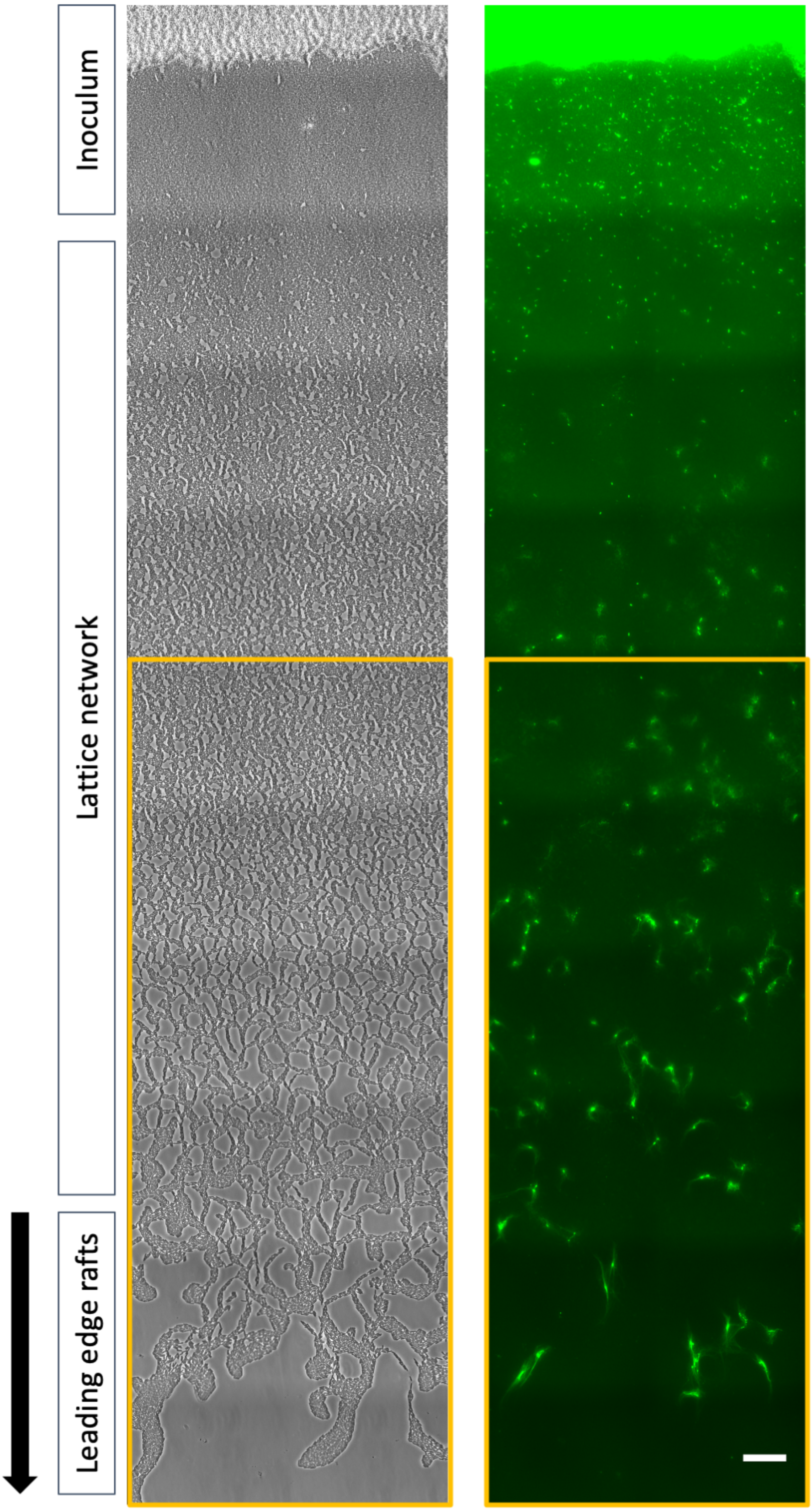

**Figure S1. Visualisation and analysis of eDNA release in actively expanding interstitial biofilms.** Representative (n=30) stitched image of a PAO1 interstitial biofilm after incubation for 4 h at 37°C. Phase-contrast (left) or EtHD-2 stained eDNA (green, right). The direction of expansion is indicated by the arrow. The orange boxed area corresponds to the 148.75 μm x 400.80 μm (59.6 $mm^2$) area analysed in Figure 1C. Scale bar=20 μm. The amount of EtHD-2 staining in the inoculum was similar to this image across all strains and images analysed.

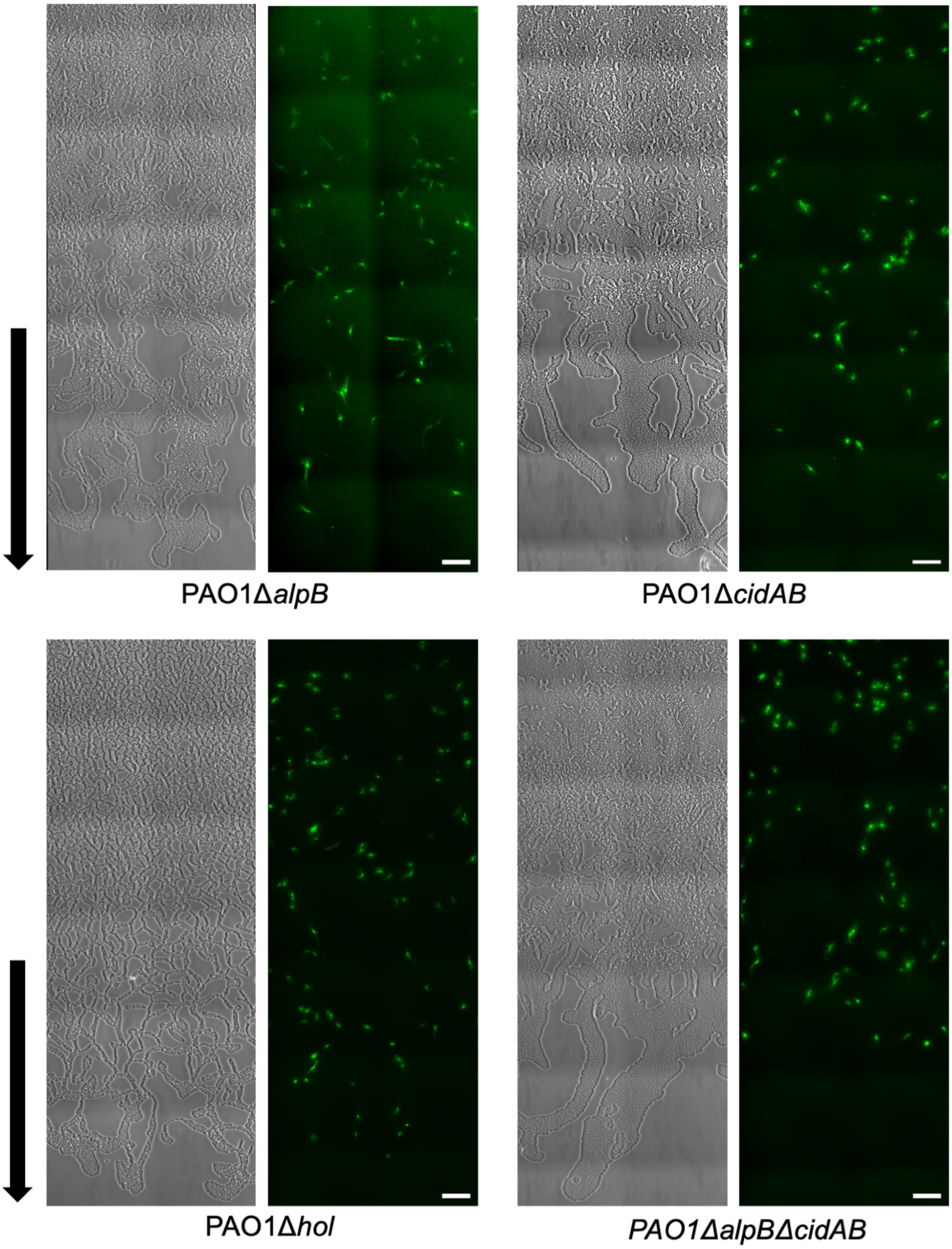

**Figure S2. Visualisation and analysis of eDNA release in actively expanding interstitial biofilms.** (Continued next page)

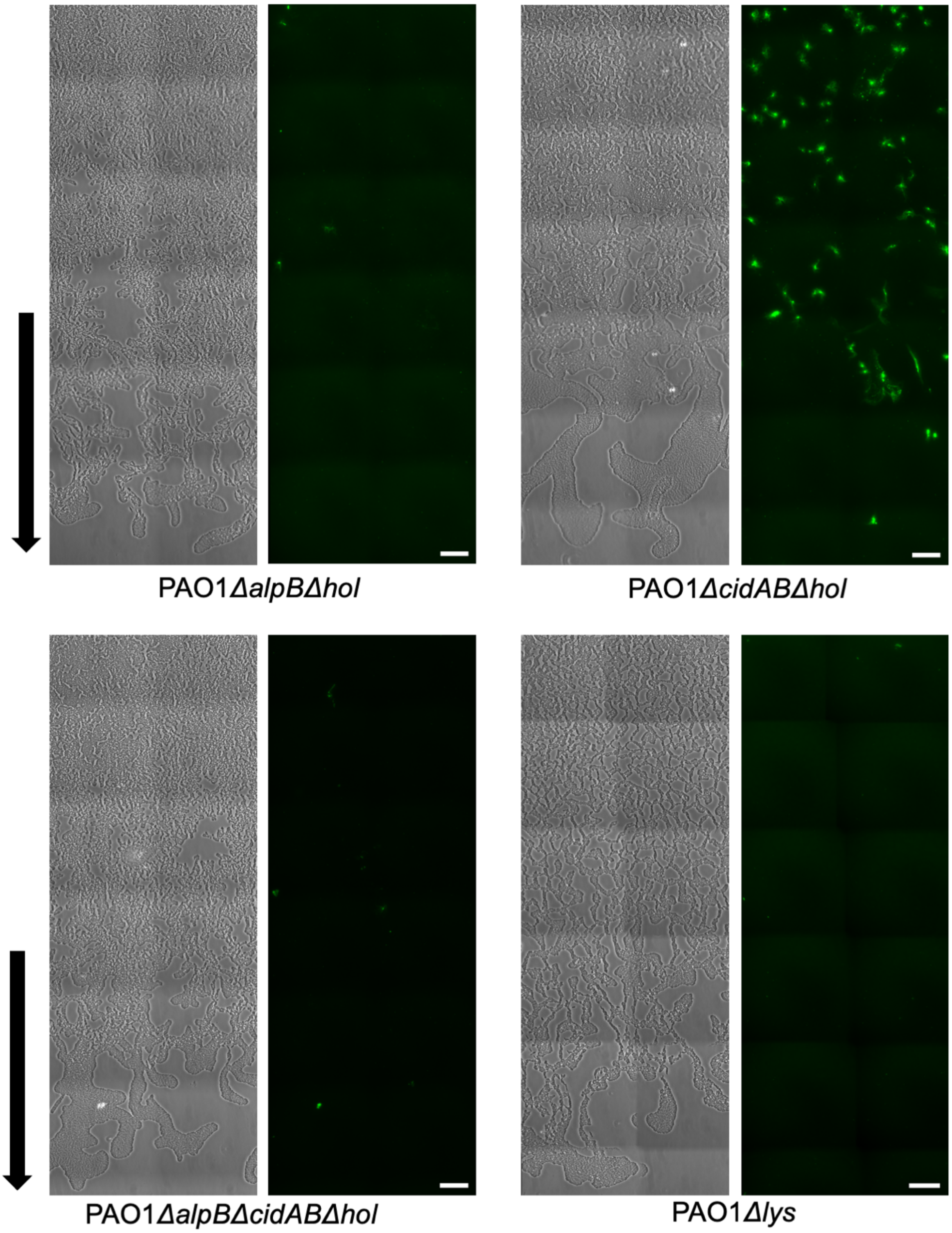

**Figure S2. Visualisation and analysis of eDNA release in actively expanding interstitial biofilms.** Representative (n=30) stitched images of PAO1Δ*alpB,* PAO1Δ*cidAB,* PAO1Δ*hol, PAO1ΔalpBΔcidAB, PAO1ΔalpBΔhol, PAO1ΔcidABΔhol, PAO1ΔalpBΔcidABΔhol* or PAO1Δ*lys* interstitial biofilms after incubation for 4 h at 37°C. Phase-contrast (left) or EtHD-2 stained eDNA (green, right). The direction of expansion is indicated by the arrow. The area visualised here is the same as the orange boxed area (48.75 µm x 400.80 µm (59.6 $mm^2$)) shown in Figure S1. Scale bar=20 µm.

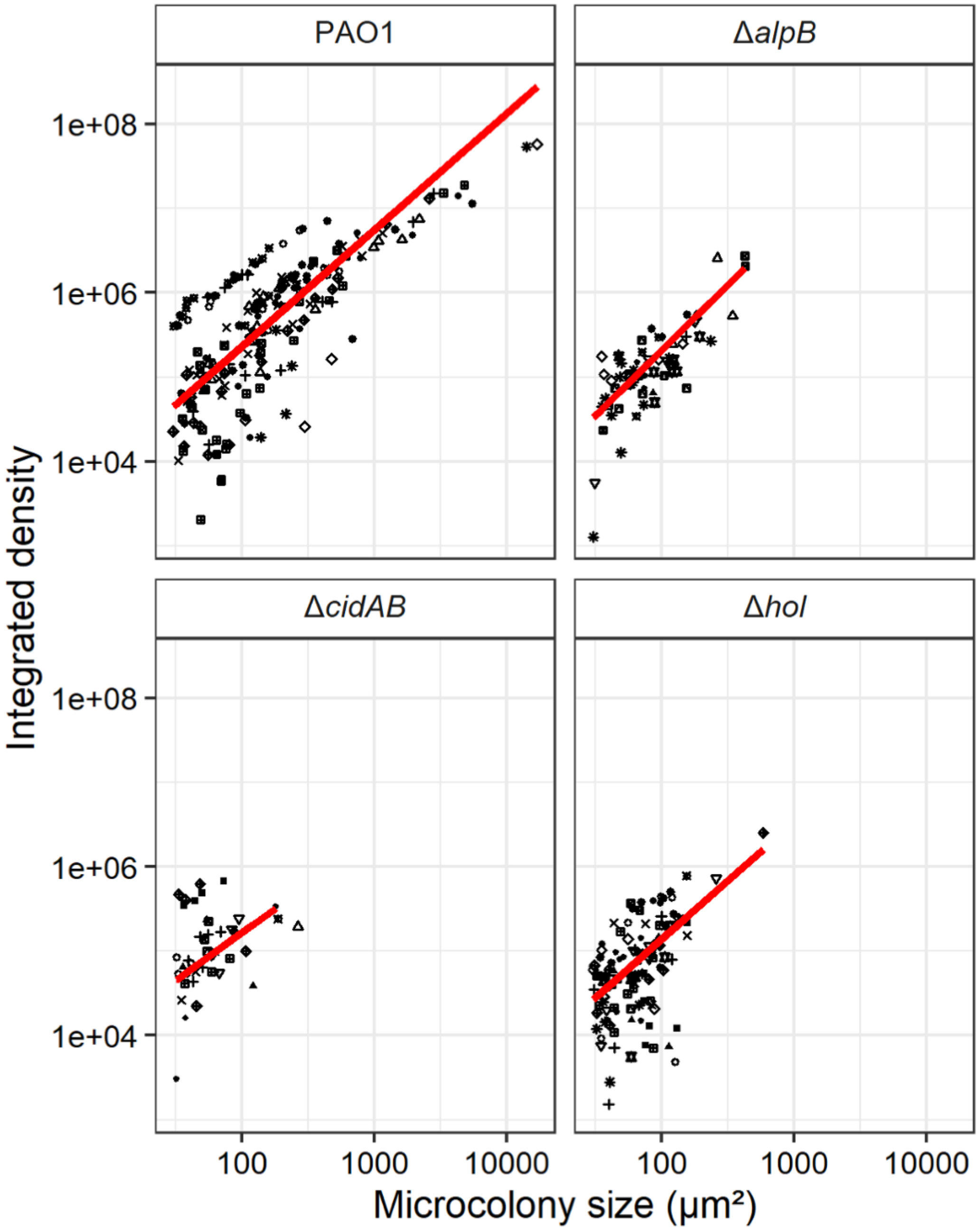

**Figure S3. Microcolony size correlates with amount of eDNA present.** The relationship between microcolony size and integrated density of fluorescent signal (eDNA) is presented using data from Figure 2C-D. In this figure shapes indicate individual fields of view. Lines are estimated slopes from mixed effects regression of log size on log density and the interaction of density with genotype. Field of view is included as a random effect. Genotype does not significantly affect the relationship between density and size (p=0.2558 compared to a model excluding the interaction term) or integrated density conditional on size (p=0.6795 compared to a model excluding the effect of genotype on density).

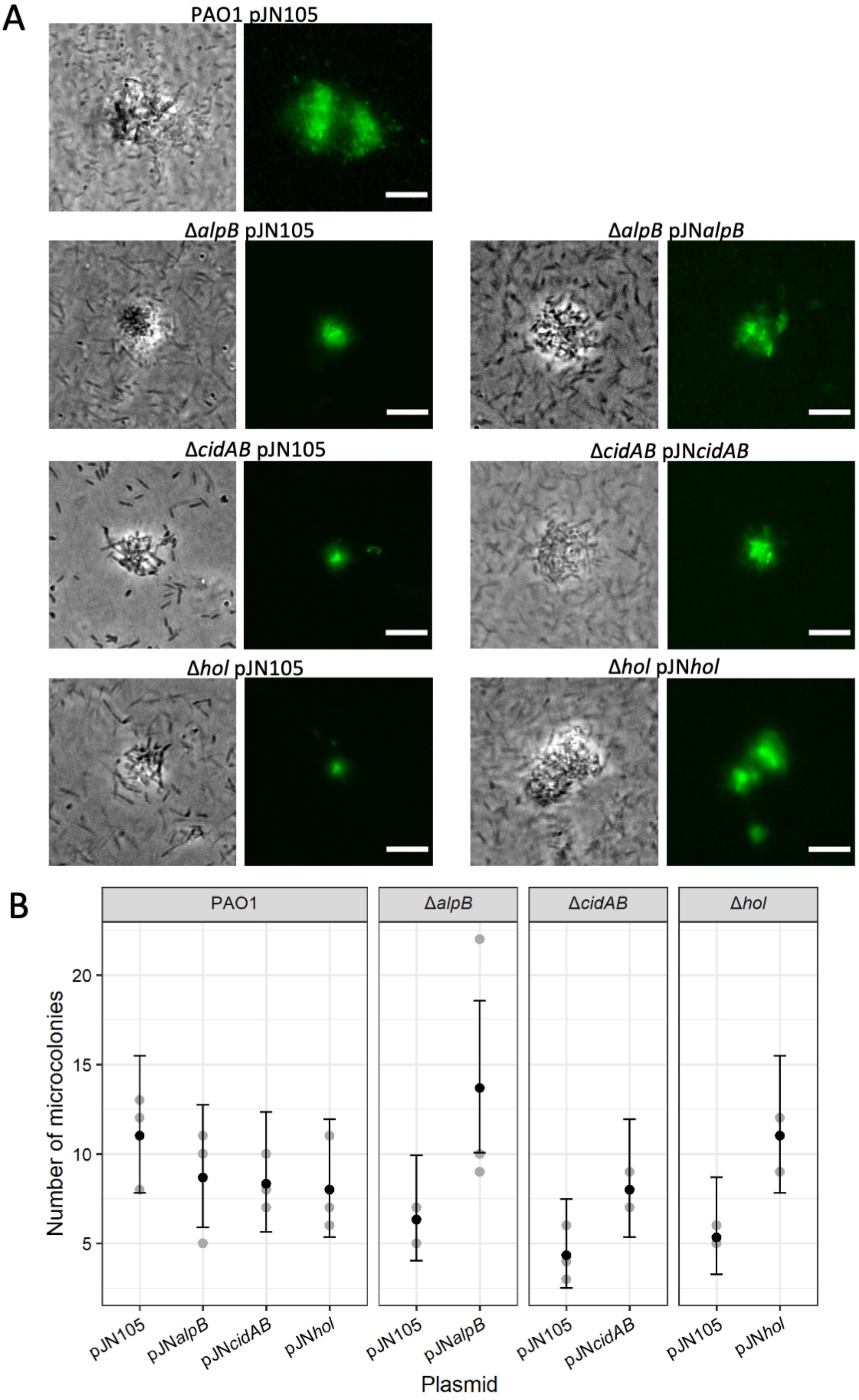
A
PAO1 pJN105
ΔalpB pJN105
ΔalpB pJNalpB
ΔcidAB pJN105
ΔcidAB pJNcidAB
Δhol pJN105
Δhol pJNhol
B
PAO1
ΔalpB
ΔcidAB
Δhol
Number of microcolonies
20
15
10
5
pJN105
pJNalpB
pJNcidAB
pJNhol
pJN105
pJNalpB
pJN105
pJNcidAB
pJN105
pJNhol
Plasmid

**Figure S4. Complementation restores microcolony formation in single holin deletions.** Microcolonies formed after 8 h incubation at 37°C for PAO1, PAO1Δ*alpB,* PAO1Δ*cidAB* and PAO1Δ*hol* with empty pJN105 plasmd, or the complementation plasmid pJN*alpB*, pJN*cidAB* or pJN*hol* induced with 0.02% L-arabinose. (A) Representative images of each strain with phase contrast (left) and eDNA (EthHD-2, right). Scale bar=10 μm. PAO1 with pJN*alpB*, pJN*cidAB* or pJN*hol* were indistinguishable from PAO1 with pJN105. (B) Total number of microcolonies formed in 8 h submerged biofilms. Estimates of means and 95% CIs were calculated using Poisson regression. See Table S7 for estimates of rate ratios for estimates, 95% CIs and p-values corresponding to the effect of plasmids, deletions, and their interactions.

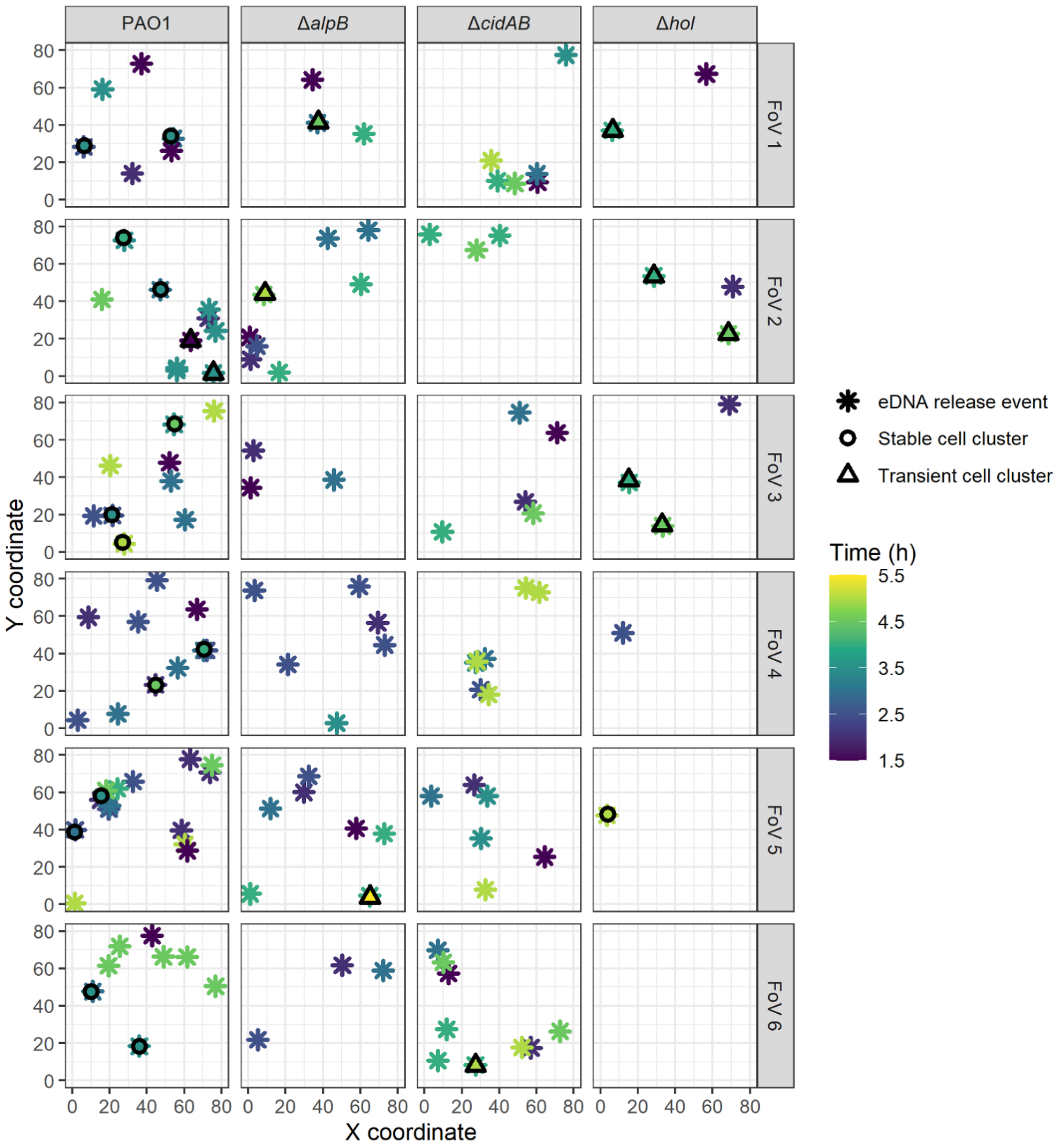

**Figure S5-S6: eDNA release events initiate the formation of cell clusters in submerged biofilms.** (Continued next page)

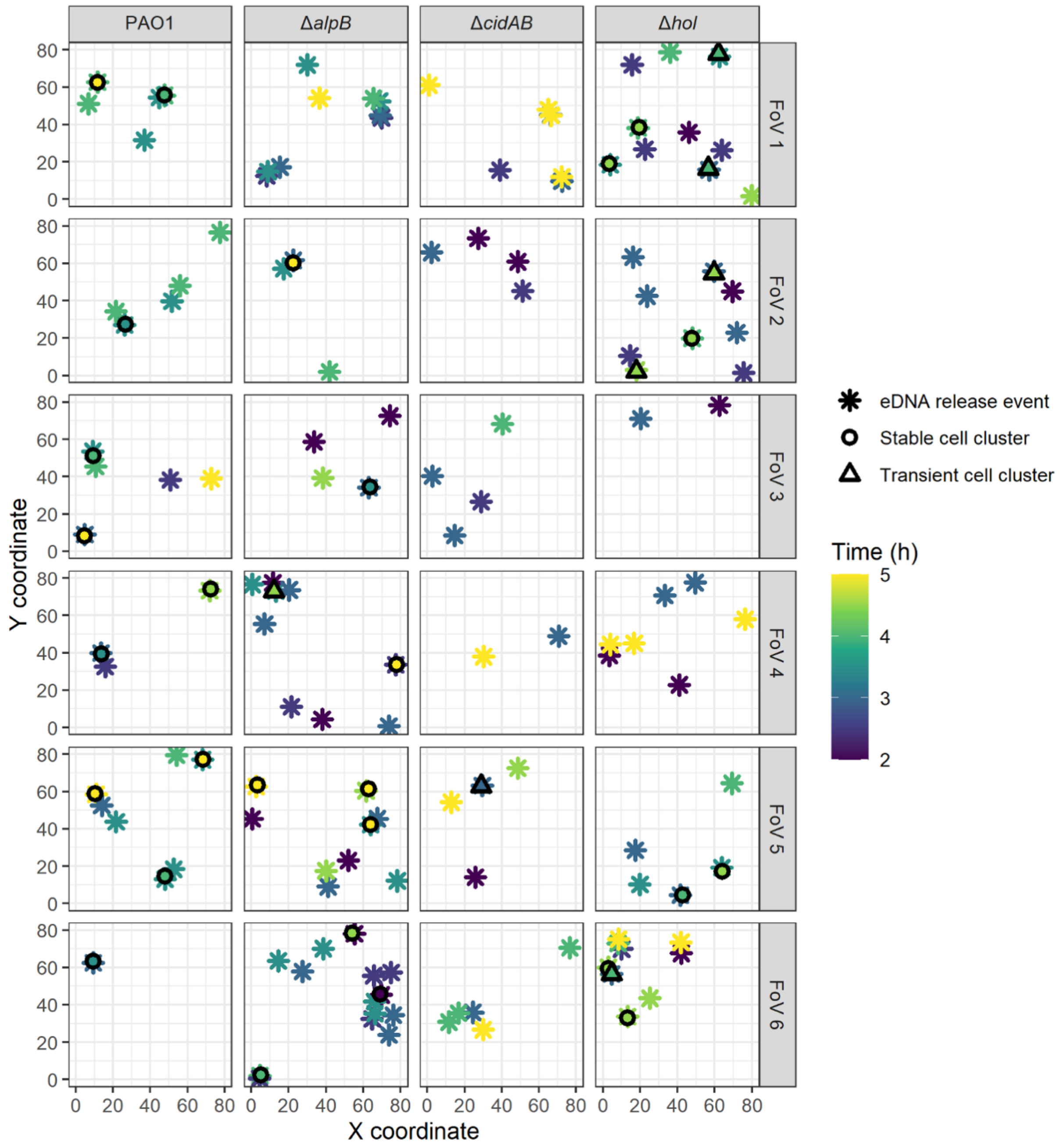

**Figure S5-S6: eDNA release events initiate the formation of cell clusters in submerged biofilms.** Cell clusters (stable or transient) and eDNA (as visualised with TOTO-1 stain) release events by PAO1, PAO1Δ*alpB,* PAO1Δ*cidAB* and PAO1Δ*hol* from 1-5.5 h at 37°C represented as X, Y coordinates for eDNA release events and associated stable or transient cell clusters. Data are combined from 12 random fields of view (FoV) across biological replicate number 1 (Figure S5) or number 2 (Figure S6) (n=12).

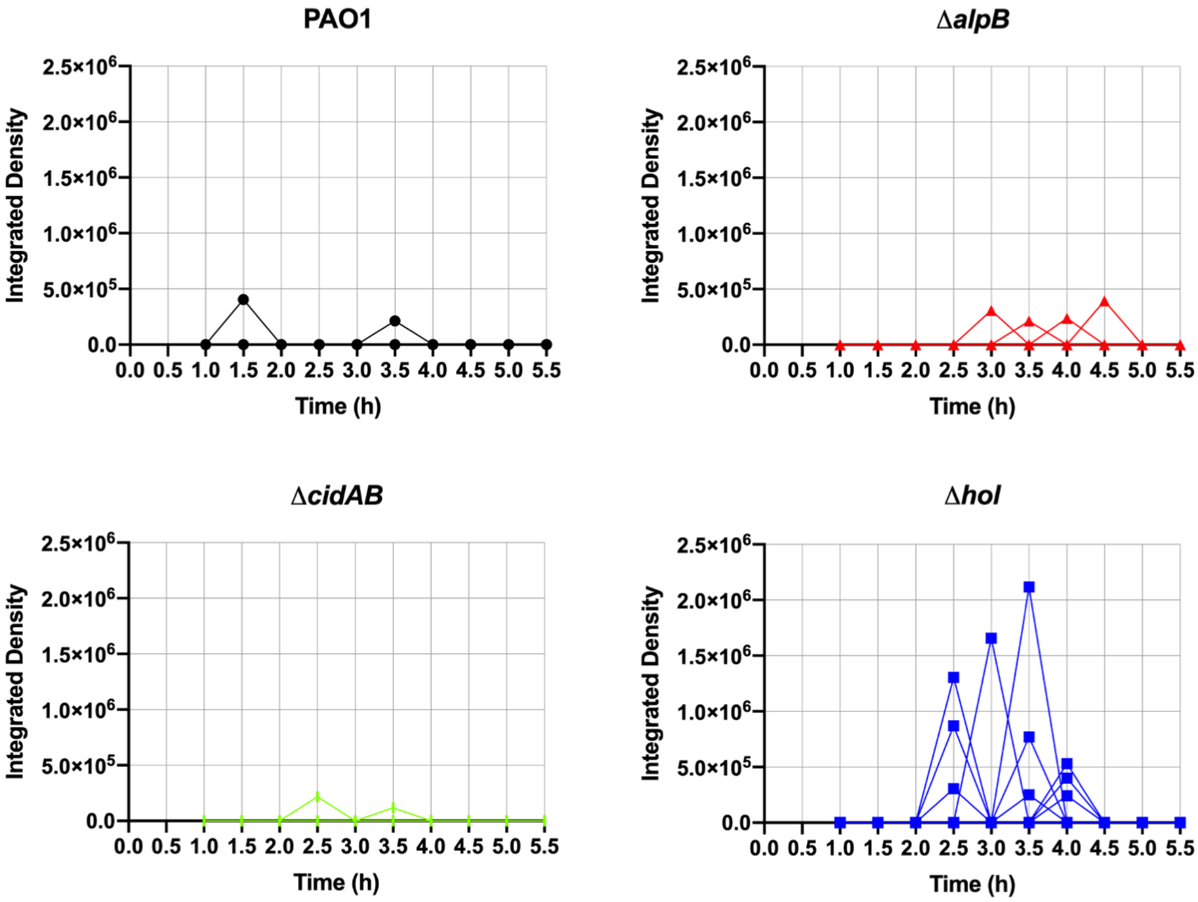

**Figure S7. Integrated density of eDNA over time in transient cell clusters.** Integrated density of fluorescence signal (eDNA – as visualised with TOTO-1 stain) associated with transient cell clusters formed over 1-5.5 h for PAO1 (2 clusters), PAO1Δ*alpB* (4 clusters), PAO1Δ*cidAB* (2 clusters) or PAO1Δ*hol* (10 clusters). Original data was generated from analysis of 12 random fields of view across 2 biological replicates (n=12).

### References

1. Simon R, O’Connell M, Labes M, Pühler A. Plasmid vectors for the genetic analysis and manipulation of rhizobia and other gram-negative bacteria. Plant Molecular Biology. Elsevier; 1986. p. 640–59.
2. Ma L, Conover M, Lu H, Parsek MR, Bayles K, Wozniak DJ. Assembly and development of the *Pseudomonas aeruginosa* biofilm matrix. PLoS Pathog. 2009 Mar 27;5(3):e1000354.
3. Turnbull L, Toyofuku M, Hynen AL, Kurosawa M, Pessi G, Petty NK, et al. Explosive cell lysis as a mechanism for the biogenesis of bacterial membrane vesicles and biofilms. Nat Commun. 2016 Apr 14;7:11220.
4. Newman JR, Fuqua C. Broad-host-range expression vectors that carry the L-arabinose-inducible *Escherichia coli* araBAD promoter and the araC regulator. Gene. 1999 Feb 18;227(2):197–203.
5. Alm RA, Mattick JS. Identification of two genes with prepilin-like leader sequences involved in type 4 fimbrial biogenesis in *Pseudomonas aeruginosa*. J Bacteriol. 1996 Jul;178(13):3809–17.
6. Hoang TT, Karkhoff-Schweizer RR, Kutchma AJ, Schweizer HP. A broad-host-range Flp-FRT recombination system for site-specific excision of chromosomally-located DNA sequences: application for isolation of unmarked *Pseudomonas aeruginosa* mutants. Gene. 1998 May 28;212(1):77–86.
